# Human-mouse proteomics reveals the shared pathways in Alzheimer’s disease and delayed protein turnover in the amyloidome

**DOI:** 10.1101/2024.10.25.620263

**Authors:** Jay M. Yarbro, Xian Han, Abhijit Dasgupta, Ka Yang, Danting Liu, Him K. Shrestha, Masihuz Zaman, Zhen Wang, Kaiwen Yu, Dong Geun Lee, David Vanderwall, Mingming Niu, Huan Sun, Boer Xie, Ping-Chung Chen, Yun Jiao, Xue Zhang, Zhiping Wu, Yingxue Fu, Yuxin Li, Zuo-Fei Yuan, Xusheng Wang, Suresh Poudel, Barbora Vagnerova, Qianying He, Andrew Tang, Patrick T. Ronaldson, Rui Chang, Gang Yu, Yansheng Liu, Junmin Peng

## Abstract

Murine models of Alzheimer’s disease (AD) are crucial for elucidating disease mechanisms but have limitations in fully representing AD molecular complexities. We comprehensively profiled age-dependent brain proteome and phosphoproteome (*n* > 10,000 for both) across multiple mouse models of amyloidosis. We identified shared pathways by integrating with human metadata, and prioritized novel components by multi-omics analysis. Collectively, two commonly used models (5xFAD and APP-KI) replicate 30% of the human protein alterations; additional genetic incorporation of tau and splicing pathologies increases this similarity to 42%. We dissected the proteome-transcriptome inconsistency in AD and 5xFAD mouse brains, revealing that inconsistent proteins are enriched within amyloid plaque microenvironment (amyloidome). Determining the 5xFAD proteome turnover demonstrates that amyloid formation delays the degradation of amyloidome components, including Aβ-binding proteins and autophagy/lysosomal proteins. Our proteomic strategy defines shared AD pathways, identify potential new targets, and underscores that protein turnover contributes to proteome-transcriptome discrepancies during AD progression.

---

Alzheimer’s disease (AD), a progressive neurodegenerative disorder, is the most common cause of dementia, affecting more than 6 million Americans^1^. AD pathology initiates decades before the onset of gross behavioral symptoms and is primarily defined by the aggregation of β-amyloid peptide (Aβ) in extracellular plaques and of hyperphosphorylated tau proteins as intracellular neurofibrillary tangles^2–4^. In addition to Aβ and tau, other coexisting molecular changes^4,5^, such as α-synuclein^6,7^, TDP-43^5,8^, and U1 snRNP^9,10^, may play important roles in disease progression. Genetic analyses of AD and control cases have elucidated three causative genes (*APP*, *PSEN1*, and *PSEN2*), high-risk genes (*APOE4* and *TREM2*) and about 100 low-risk genes and loci^11–20^, However, the molecular mechanisms of these proteins/genes in AD development are not fully understood, often due to the lack of suitable cellular or animal models.

More than 100 genetic AD mouse models have been developed^21–24^, predominantly by mimicking genetic mutations linked to familial AD, such as the lines of 5xFAD^25,26^, 3xTG^27^, and APP-KI including APP^NLF^ (NLF) and APP^NLGF^ (NLGF)^28^. However, none of these models capture the full spectrum of AD molecular events and pathologies as they exhibit less severe neurodegeneration compared to human patients. Ideally, researchers should fully understand the advantages and limitations of mouse models to select the most appropriate one for addressing specific hypotheses; however, no comprehensive resources are currently available.

Rapid developments in omics technologies provide an opportunity of thoroughly evaluating disease models on a global scale and exploring their relevance by comparisons with human data^29–31^. Transcriptomic analyses of the amyloidosis mouse models revealed changes in expression of genes linked to immune response, synaptic function, and neuronal signaling^32–34^. However, RNA levels do not always align with protein levels due to posttranscriptional processes, such as translation and protein turnover^35^. Indeed, notable inconsistencies between transcript and protein levels in AD and 5xFAD mice were observed^36,37^. Complementary proteomic studies in AD mice^38–41^ not only corroborated transcriptomic findings, but also identified RNA-independent protein alterations^36,39,42^ and changes in protein turnover^43,44^. These early proteomic studies in the AD mice uncovered some molecular changes but they were often limited by inadequate proteomic depth, restricted analysis of individual mouse models, and/or insufficient comparison with human AD datasets.

Here we present a deep, age-dependent analysis of 10,369 proteins (10,331 genes) and 12,096 phosphopeptides (10,532 phosphosites) across commonly used AD models: 5xFAD^25^, NLF^28^, and NLGF^28^. We also profiled two additional AD models (3xTG^27^ and BiG^45^), performed human-mouse comparisons, and analyzed transcriptome-proteome inconsistency in both mouse and human. To explore the contribution of protein degradation to the transcriptome-proteome inconsistency, we measured the turnover rates of 8,492 brain proteins and found that amyloid formation delays the degradation of amyloidome components. Thus, our comprehensive proteomic analysis identifies shared AD pathways and demonstrates altered protein turnover in amyloid plaques in AD mice. All data are freely available and searchable through an interactive website (https://penglab.shinyapps.io/mouse_ad_profile/).

## Results

### Proteome profiling of multiple AD amyloidosis models reveals shared pathways

We compared proteomic readout of three mouse models of amyloidosis at different ages, with early and late symptoms (**Fig. 1a**, **Supplementary Table 1**): (i) 5xFAD (3-, 6-, 12-month-old), overexpressing human *APP* and *PSEN1* genes carrying a total of five human disease mutations under the *Thy1* promoter, which promotes rapid onset of amyloid pathology^25^; (ii) NLF (3-, 12-month-old, with weak pathology) and NLGF (3-, 6-, 12-, 18-month-old, with strong pathology), both being next-generation knock-in models with humanized Aβ without gene overexpression^28^. We also analyzed age-matched wild type (WT) control mice for each mouse line.

**Fig. 1.**
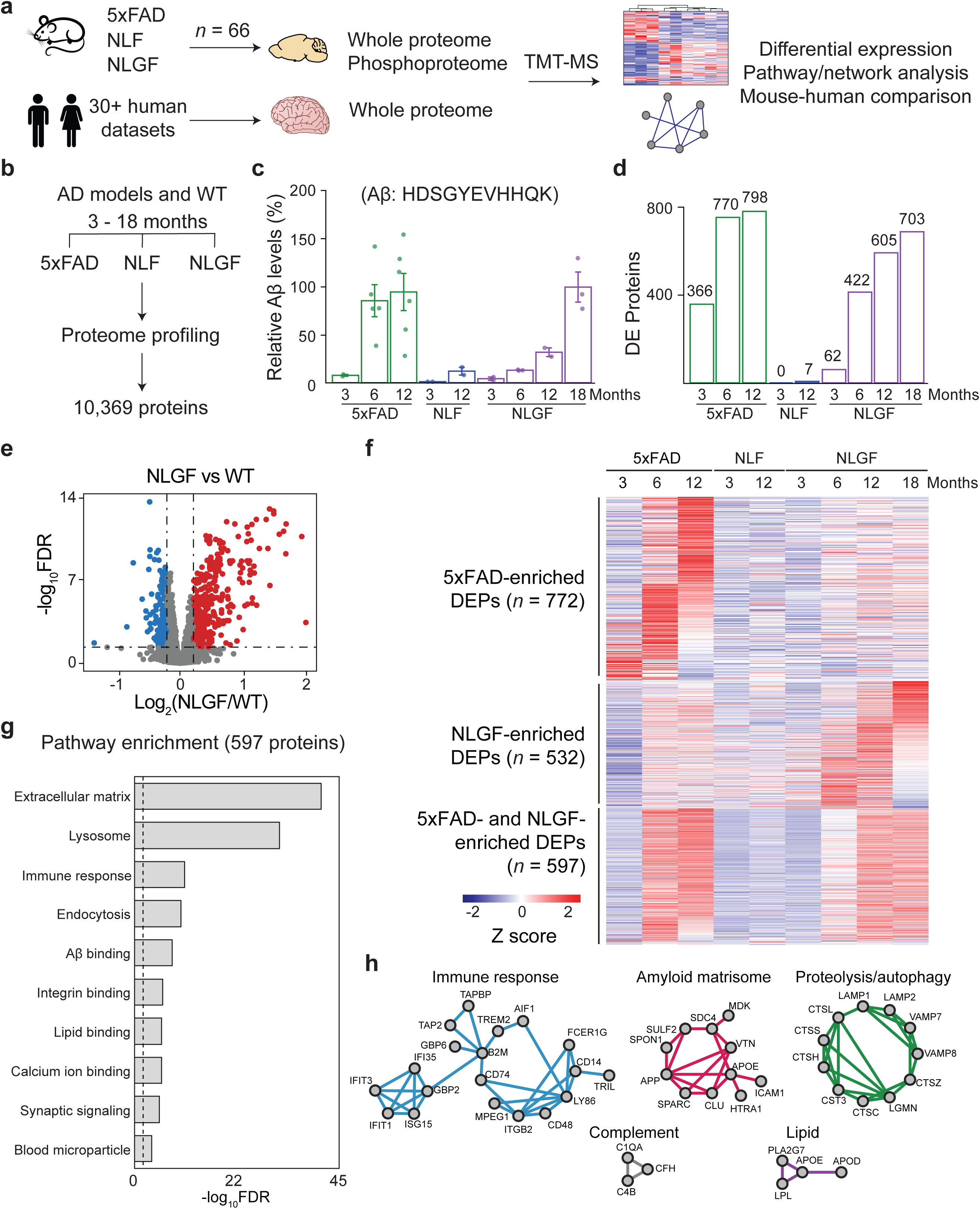
Brain proteomics reveals proteomic changes that are shared in AD mouse models. **a**, Schematic plan of this study. Mouse cortical tissues from AD models of amyloidosis (5xFAD, NLF, NLGF, and matched WT, total *n* = 66 for 16 conditions, averaged *n* = ∼4 per condition) were analyzed by TMT-LC/LC-MS/MS and compared with human metadata. **b**, Proteins quantified at different ages (3-18 months). **c**, Aβ levels quantified by MS. The values were averaged for each age and model, then normalized to 12-month-old 5xFAD (100%). **d**, DEPs between AD mice and WT controls, defined by moderated *t*-test with statistical cutoffs. **e**, Volcano plot for NLGF-WT comparison (FDR < 0.05, |log2FC| > 2SD, dashed lines). **f**, Heatmap of DEPs in AD mice, including the proteins enriched in 5xFAD or NLGF and those shared by both mice. **g**, Pathway analysis of shared DEPs in 5xFAD and NLGF. FDR was derived from *p* values (Fisher’s exact test) by the Benjamini-Hochberg procedure. **h**, Enriched PPI modules from biological processes using the shared DEPs.

Using our optimized tandem mass tag (TMT) method, coupled with extensive two-dimensional liquid chromatography (LC/LC) and high-resolution tandem mass spectrometry (MS/MS, **Extended Data Fig. 1a**)^46–49^, we profiled a total of 66 mouse brains (cortex) in multiple TMT batches with deep proteome coverage (**Supplementary Table 2**), identified more than 900,000 peptide-spectrum matches (PSMs), ∼330,000 peptides, and 10,369 unique proteins (10,331 genes) that were shared in all animals, with a protein false discovery rate (FDR) below 0.01 (**Fig. 1b, Supplementary Table 3**). After protein quantification based on TMT reporter ions, sample loading bias was corrected as shown in a box plot (**Extended Data Fig. 1b**), and the batch effect was normalized and confirmed by PCA analysis (**Extended Data Fig. 1c**). As expected, the Aβ tryptic peptide (R.HDSGYEVHHQK.L) shows age-dependent increases in all AD mice, with higher levels in 5xFAD and NLGF than NLF, consistent with the reported pathologies in these mice (**Fig. 1c, Supplementary Table 4**).

We examined the effect of aging using only WT mice (3-, 6-, 12-, 18-month-old) to avoid the confounding impact of Aβ insult in different genotypes. When comparing 3-month-old mice to any other aged mice, differential expression (DE) analysis identified 183 age-dependent proteins (FDR < 0.05, |log_2_FC| > two standard deviation (SD)), with 129 proteins upregulated and 54 proteins downregulated with age (**Extended Data Fig. 2a**, **Supplementary Table 5**). The upregulated proteins are enriched in the Gene Ontology (GO)^50^ terms of collagen-containing extracellular matrix, perineuronal net, lysosome, glutathione metabolic process, etc. Conversely, the downregulated proteins are enriched in cell periphery, cell junction, and neuronal components, including synapse, axon, dendritic tree, dendritic spine, etc. (**Extended Data Fig. 2b-c, Supplementary Table 6**).

We then performed DE analysis at different ages for each genotype using WT controls (FDR < 0.05, |log_2_FC| > 2SD, **Supplementary Table 7**), excluding human Aβ peptide as it is not present in the WT mice. NLF exhibits a few DE proteins (DEPs), in agreement with its weak pathology^51^ (**Fig. 1d**). In contrast, the 5xFAD and NLGF models demonstrate significant protein alterations, which increased with age for both models (**Fig. 1d**). For instance, a volcano curve shows 605 DEPs in 12-month-old NLGF mice compared to the WT (**Fig. 1e**), reflecting its strong amyloid phenotype^25,28^. When summing the DEPs at different ages, there are 1,382 DEPs in 5xFAD, only 7 in NLF and 1,142 in NLGF. Moreover, we found that the numbers of DEPs are highly correlated with the Aβ accumulation across all tested models (R = 0.86, **Supplementary Table 8**).

In spite of the genetic difference between 5xFAD and NLGF models^21,28^, we observed comparable proteomic signatures in the two models. Among 1,914 total DEPs in 5xFAD and NLGF, 610 (32%) overlap, of which 98% (597/610) change in the same direction. The remaining DEPs, 772 (40%) in 5xFAD and 532 (28%) in NLGF, are genotype-specific, but show a similar age-dependent trend in both AD models (**Fig. 1f**).

We performed Gene Ontology analysis on the 597 overlapping DEPs and found enrichment in extracellular matrix, lysosome/endocytosis, immune response, synaptic signaling, and binding to Aβ, integrin, lipid, calcium ion, etc. (**Fig. 1g, Supplementary Table 9**). The overlapping DEPs were mapped to protein-protein interaction (PPI) networks, revealing significant interacting modules of immune response, complement, lipid metabolism, proteolysis/autophagy, and an amyloid-binding extracellular matrix protein network (amyloid matrisome)^37^ (**Fig. 1h**). These findings suggest that proteins and pathways shared between 5xFAD and NLGF are critical for amyloidosis pathogenesis and immune response, consistent with previous reports in human AD^36,37,52^.

### Phosphoproteome profiling of AD models highlights alterations independent of protein levels

Since protein phosphorylation is known to contribute to AD pathogenesis^53,54^, we profiled the phosphoproteome in the same set of 5xFAD (12-month-old) and NLGF (3-, 6-, 12-, 18-month-old) mice and their age-matched WT controls. The profiling was also performed using the TMT-LC/LC-MS/MS method^46–49^, with an additional step of phosphopeptide enrichment to improve the phosphoproteome coverage^55^ (**Extended Data Fig. 3a-c**). When combining phosphoproteome data from all TMT batches, we quantified 129,906 unique phosphopeptides (82,261 phosphosites on 10,502 proteins, peptide FDR < 0.01, **Supplementary Table 10**). Because of the incomplete coverage of phosphoproteome in individual TMT batches, the overlap between different TMT batches is often low^56^. When we extracted the phosphopeptides shared in all 36 mice, the number dropped to 12,096 phosphopeptides (10,532 phosphosites on 2,814 proteins, **Fig. 2a, Supplementary Table 11**). These shared phosphopeptides contain 8,665 pS (82.3%), 1,625 pT (15.4%), and 242 pY (2.3%) sites (**Fig. 2b**), similar to the site distribution in other large phosphoproteome analyses^36,57^.

**Fig. 2.**
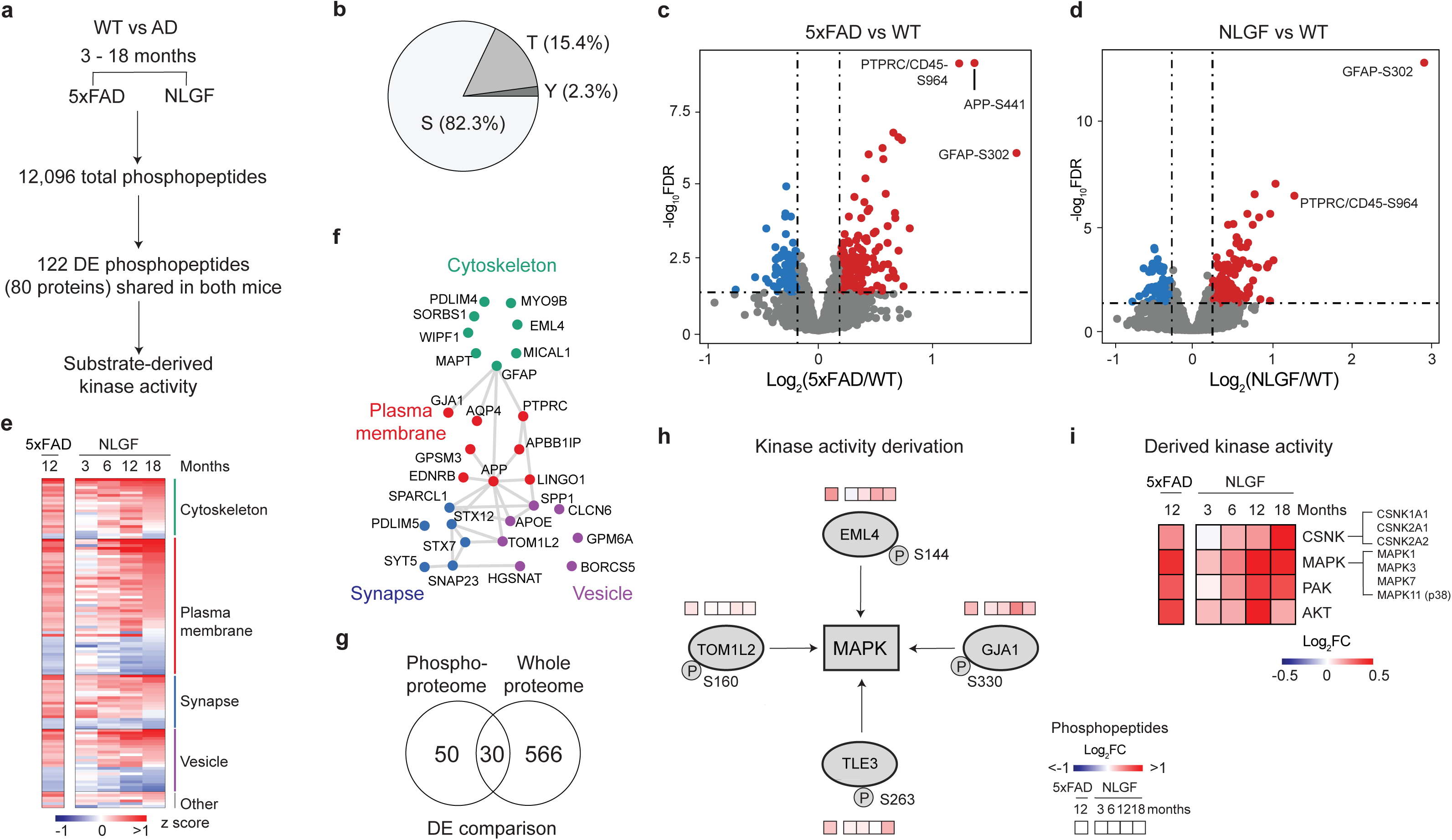
Tissue phosphoproteomics defines a new layer of regulation beyond whole proteome. **a**, Phosphoproteome profiling was performed across 36 5xFAD and NLGF mice and their age-matched controls. We quantified 12,096 phosphopeptides (peptide FDR < 0.01) shared in all mice and performed statistical comparisons by moderated *t*-test. 122 DE phosphopeptides (80 proteins) were identified (FDR < 0.05, |log_2_FC| > 2SD). **b**, Distribution of phospho-Ser/Thr/Tyr in identified phosphosites. **c**, Volcano plot of phosphoproteome data for 12-month-old 5xFAD compared to WT. Dashed lines indicate cutoffs. **d**, Volcano plot of phosphoproteome data for 12-month-old NLGF compared to WT. **e**, Heatmap of DE phosphoproteins in 5xFAD and NLGF mice, with protein subcellular location shown. **f**, PPI modules of DE phosphoproteins. **g**, The overlap of DEPs in phosphoproteomics and whole proteome analysis. Only consistent DEPs in both AD models were counted. **h**, Bioinformatics method for identifying altered kinase activities. Kinase-substrate linkages were extracted to infer kinase activities by the KSEA algorithm. An examples of MAPK activation based on KSEA. Phosphopeptide levels are displayed using the accompanying gradients. **i**, Heatmap of derived kinase activities. The fold change (FC) of 5xFAD and NLGF was calculated by comparison with WT.

We then carried out pairwise DE analysis using age-matched WT controls for both 5xFAD and NLGF mice, identifying 122 consistent DE phosphopeptides (80 DE phosphoproteins) in the two mouse models (FDR < 0.05, |log_2_FC| > 2SD, **Supplementary Table 12**). For example, in both mice, the phosphorylation levels of PTPRC/CD45 S964 and GFAP T299 are significantly increased, indicating the activation of microglia and astrocytes, respectively (**Fig. 2c and 2d**). The 80 DE phosphoproteins are enriched in several pathways and PPI networks of cytoskeleton, plasma membrane, synapse, and vesicle (**Fig. 2e and 2f, Supplementary Table 13**). These pathways are consistent with those revealed in phosphoproteomic studies of human AD^54,58^.

Notably, only 30 (37.5%) of the 80 DE phosphoproteins showed significant changes at the proteome level (**Fig. 2g**), suggesting that the differences in phosphorylation are primarily due to altered kinase/phosphatase activity in the brain, independent of the protein levels. To quantify the change in kinase activities based on these DE phosphopeptides, we derived alterations in kinase families by the computer algorithm of kinase-substrate enrichment analysis (KSEA)^59^. For instance, the MAPK activity could be inferred from its DE substrates of TOM1L2, EML4, GJA1 and TLE3 (**Fig. 2h**). The analysis identified the upregulation of four kinase families: casein kinase (CSNK), mitogen-activated protein kinase (MAPK, including p38 kinase), p21-activated kinase (PAK), and protein kinase B (AKT) (**Fig. 2i**). Consistently, MAPK pathway deregulation was previously highlighted in several cohort studies of human AD^36,37,54^. Thus, our KSEA analysis suggests the deregulation of numerous kinases that are relevant to amyloid pathology.

### Human-mouse comparison identifies shared AD pathways

To investigate the relevance of the mouse models, we compared the DEPs in 5xFAD and NLGF with human metadata^36,37,52,60^. We focused on 866 human DEPs that consistently exhibited significant changes in deep AD proteomics studies (**Fig. 3a, Supplementary Table 14**), for which 654 homologous proteins were detected by MS in mice. The sum of 5xFAD and NLGF DEPs corresponds to 30% (196/654) of the AD DEPs and demonstrates age-dependency (**Fig. 3b**). The three datasets share a core set of 108 DEPs (**Fig. 3c**). Further analysis reveals that 5xFAD and NLGF DEPs align more closely with late-stage AD (R = 0.32 and 0.23, respectively) than with mild cognitive impairment (MCI) (R = 0.00 and -0.09, respectively), suggesting these mouse models are more representative of the amyloidosis in the advanced AD stages rather than that in the early, asymptomatic phase (**Fig. 3d**).

**Fig. 3.**
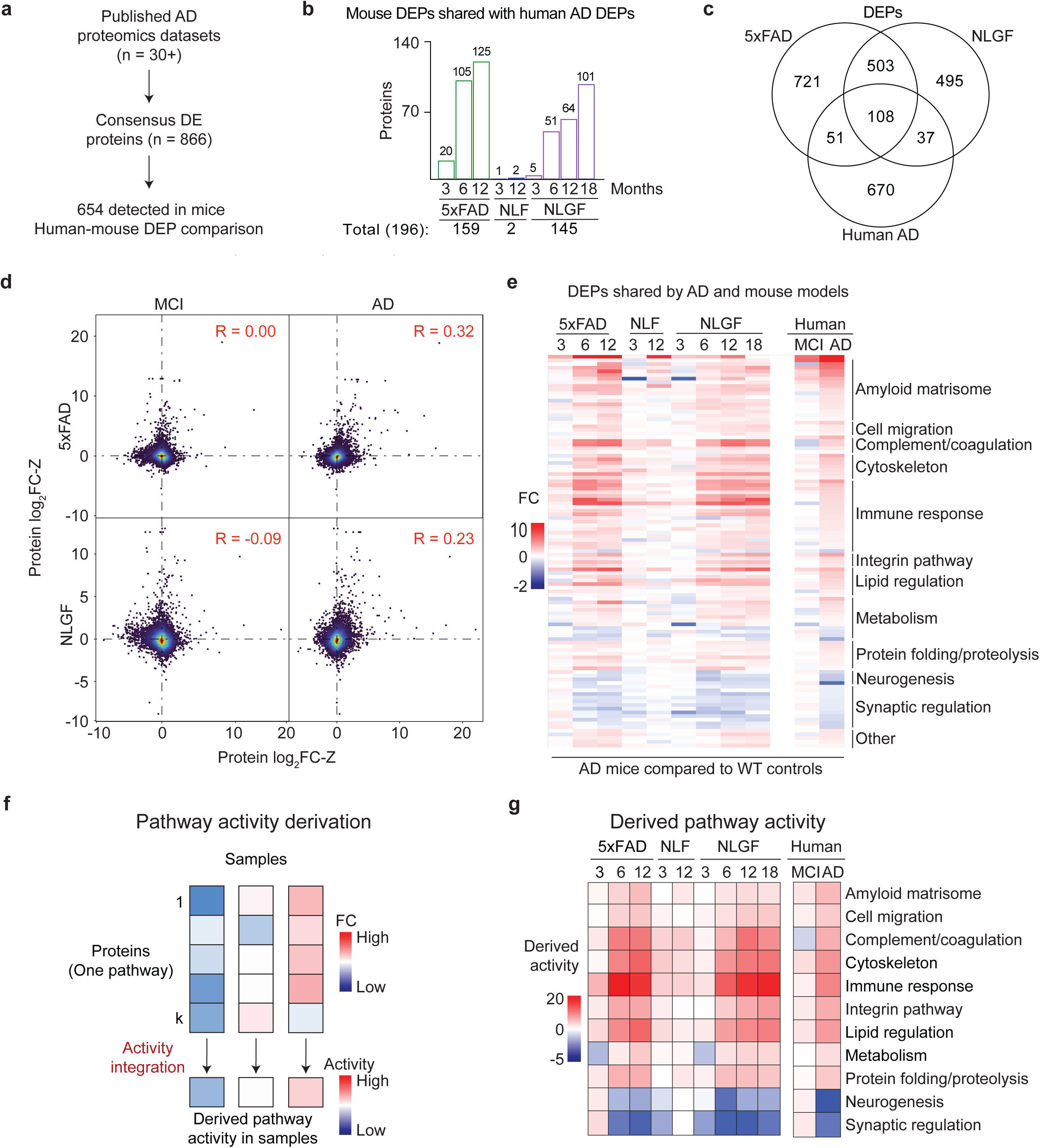
Comparison of mouse proteomics to human metadata identifies shared proteomic changes. **a**, We identified 866 proteins that are consistently altered in more than 30 human AD proteomics studies, 654 of which were quantified in the proteomic analysis of the AD mice. Of these, 196 (30%) are differentially expressed in at least one mouse model (FDR < 0.05, |log_2_FC| > 2SD). **b**, Number of overlapping DEPs between human AD and different mouse models. **c**, DEPs shared by human AD, 5xFAD, and NLGF mice. **d**, Scatter plot comparisons between Z scores of log_2_fold change values (log_2_FC-Z) of human AD/control cases and mouse models/WT at 12-month ages. Each dot represents one protein, and the color shows the dot density. Pearson correlation (R) values are shown. **e**, Heatmap showing log_2_FC values of human-mouse shared AD proteins, classified by biological pathways. **f**, Workflow for deriving pathway activities. The FC of proteins in each pathway are integrated to calculate the pathway activity. **g**, Heatmap of pathway activities in AD and mouse models.

We then examined the 108 DEPs consistent in both mouse models and human AD (**Fig. 3e**), and derived their pathway activities by integrating individual components with a mathematical formula^61,62^ (**Fig. 3f**). Several pathway activities are upregulated, including amyloid matrisome, cell migration, complement and coagulation, cytoskeleton, immune response, integrin pathway, lipid regulation, metabolism and protein folding/proteolysis; two pathways — neurogenesis and synaptic regulation — are downregulated (**Fig. 3g, Supplementary Tables 14 and 15**).

### Additional pathologies beyond amyloidosis in mice increase similarity to AD

The 5xFAD and NLGF mouse models of amyloidosis fail to capture the entire spectrum of AD-related proteomic changes. This discrepancy may be attributed to the absence of other pathologies found in human AD, such as tauopathies^53^ and splicing dysfunctions^9^. We further profiled two other AD mouse models with additional pathologies (**Fig. 4a, Extended Data Fig. 4a-e, Supplementary Table 1**): (i) 3xTG, displaying both amyloid plaque and tau tangle pathologies^27^, and (ii) BiGenic (BiG) mice, generated by crossing 5xFAD with N40K transgenic mice recapitulating both amyloid pathology and the newly discovered U1 snRNP splicing dysfunction in AD^45^. We profiled both models and age-matched controls (∼6-month-old, totaling 37 mouse brains), quantifying 9,780 and 10,255 proteins (protein FDR < 0.01). We identified 1,230 and 1,564 DEPs in 3xTG and BiG, respectively (FDR < 0.05 and |log_2_FC| > 2SD, **Fig. 4b and 4c, Supplementary Tables 16, 17**), including a substantial number of DEPs that are not observed in the pure amyloidosis models, 5xFAD and NLGF. For example, the 3xTG shows mouse-specific DEPs (**Fig. 4d**) related to tau pathways^63^, and the BiG mice uniquely displayed defective RNA splicing components^9^ and synaptic dysregulation^64^ (**Fig. 4e**).

**Fig. 4.**
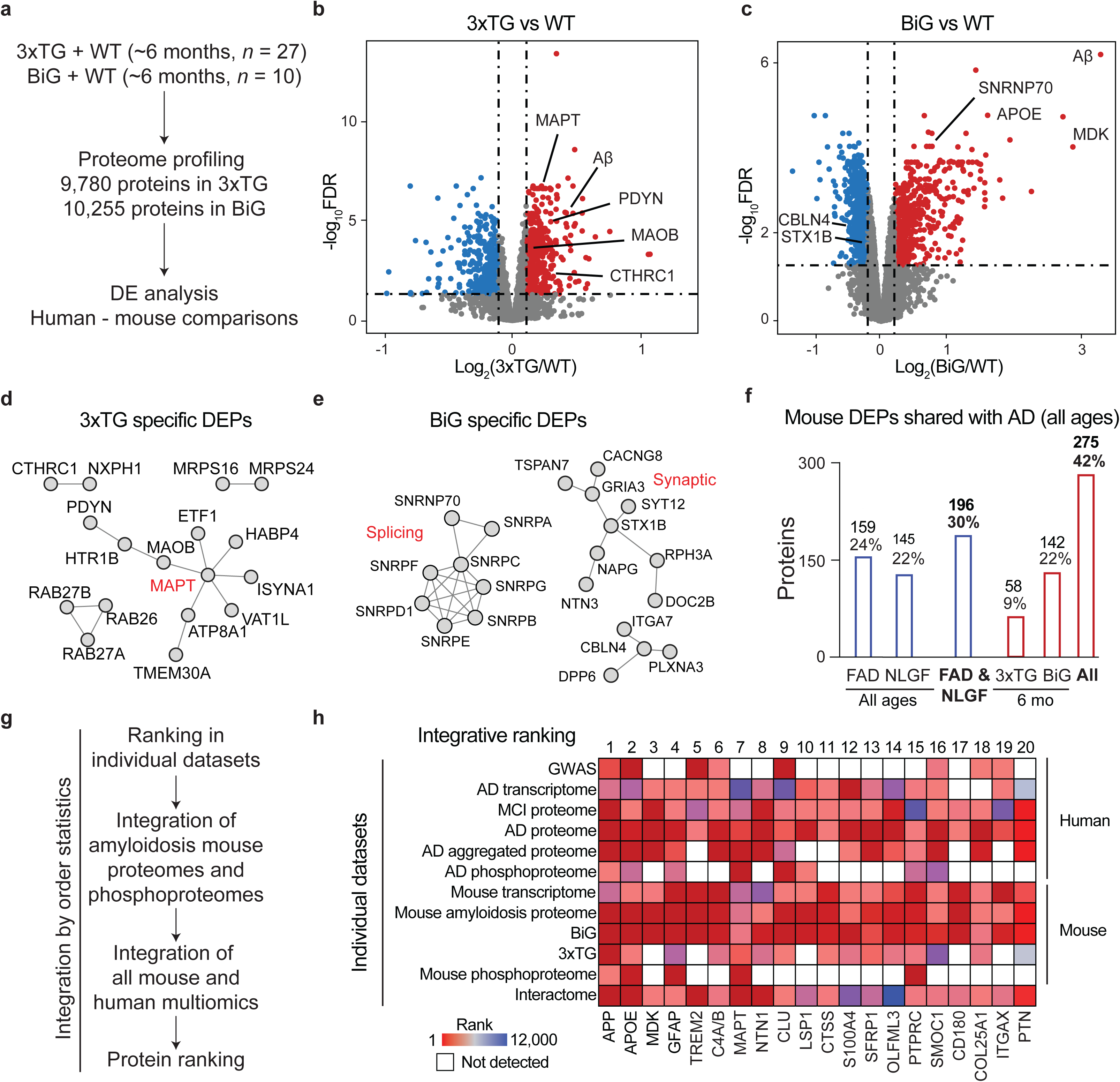
Mouse models with additional pathologies beyond amyloidosis increase the similarity to AD. **a**, Proteomic profiling of two more mouse models that express additional AD pathologies: WT (*n* = 8) and 3xTG (Aβ and tau pathologies, *n* = 19), as well as WT (*n* = 4) and BiG (Aβ and U1 splicing pathologies, *n* = 4). All mice were of ∼6-month-old. The proteomic data were subjected to DE analysis and comparison with human AD data. **b-c**, Volcano plots of log_2_(fold change) and FDR in 3xTG and BiG mice, compared to WT, with DEPs highlighted in colors and cutoffs indicated by dashed lines. **d-e**, Selected protein-protein interactions of significantly altered DEPs found exclusively in individual mice, such as MAPT interactome in 3xTG, and splicing/synaptic interactome in BiG. **f**, Numbers of DEPs in AD mouse models that were consistently altered in AD. The percentage was calculated using a denominator of 654 AD DEPs that were detectable by MS in mice. **g**, Strategy for ranking individual proteins by multi-omics using order statistics. (i) All age-dependent proteomic data from 5xFAD and NLGF were initially consolidated into two datasets for the amyloidosis proteome and phosphoproteome. (ii) These datasets were then integrated with 10 additional datasets, which include the mouse transcriptome (5xFAD), 3xTG/BiG proteomes, human genetic data from GWAS, human transcriptomes, proteomes (MCI and two independent AD studies, *n* = 3), phosphoproteome, and interactome datasets. **h**, Protein integrative rankings defined by combining 12 datasets. The entire datasets were ranked based on all identified genes/proteins. Subsequently, we extracted the rankings for the AD-mouse shared proteins (*n* = 275). The top 20 proteins are displayed, with missing values represented by white boxes.

By summing all DEPs across the four mouse models, the overlap with human DEPs increases to 42% (275/654, **Fig. 4f, Supplementary Table 14**), suggesting that additional pathologies beyond amyloid plaques contribute to alterations in the mouse proteome, moving it closer to the AD spectrum. The remaining 58% of human AD DEPs show enrichment in the pathways of mitochondrial function, cell morphogenesis, lipid regulation, potentially due to reduced neuronal cell death in the mouse models and difference in response to pathological insults between mice and human (**Supplementary Table 18**).

Among the 275 DEPs conserved between mouse models and human (**Fig. 4f**), we found that 86% are not well studied in the context of AD, with <20 AD-related publications (**Extended Data Fig. 4f**). We thus prioritized these proteins employing a method of order statistics^36^ by integrating available 12 omics datasets from both mouse and human, which include GWAS (*n* = 1), transcriptome (*n* = 2), proteome (*n* = 6), phosphoproteome (*n* = 2), and interactome (*n* = 1) (**Fig. 4g, Supplementary Table 19**). As expected, well-known proteins such as APP, APOE, GFAP, TREM2, MAPT (tau), and CLU rank highly, while other proteins in the top 20 list, such as MDK, NTN1, SFRP1, OLFML3, PTPRC/CD45, SMOC1, CD180, and PTN, remain understudied (**Fig. 4h**). These proteins require further investigation to understand their roles in the development of AD.

### Transcriptome-proteome inconsistency occurs in AD and the 5xFAD model

A transcriptome-proteome inconsistency has been reported in AD^36,37^. To explore that issue, we compared the quantitative transcriptome and proteome datasets from both human AD (*n* = 10,781) and 5xFAD mice (*n* = 8,840) relative to their control samples, after Z-score transformation (**Fig. 5a, Supplementary Table 20**). The transcriptome-proteome correlations were modest, with R values of 0.40 in human and 0.46 in mice (**Fig. 5b-c**). We identified the RNA-independent protein changes in the following steps: (i) identifying Z-score changed proteins in human AD (*n* = 1,121) and in 5xFAD mice (*n* = 1,152), compared to their controls; (ii) categorizing those changed proteins into four groups based on protein up-/down-regulation and RNA dependency/independency (**Fig. 5d**, **Supplementary Table 20**). Remarkably, in both species, approximately one-third of the altered proteins exhibit RNA independence (35% in humans and 36% in mice). We asked whether these RNA-independent protein changes are shared between AD and mouse models. From the 262 RNA-independent, upregulated proteins in human and 295 in mouse, there was an overlap of 31 proteins. In contrast, from the 133 RNA-independent, downregulated proteins in human and 120 in mouse, only one overlaps (**Fig. 5e**). These findings suggest a partial conservation of transcriptome-proteome inconsistency of upregulated proteins in the AD mouse model.

**Fig. 5.**
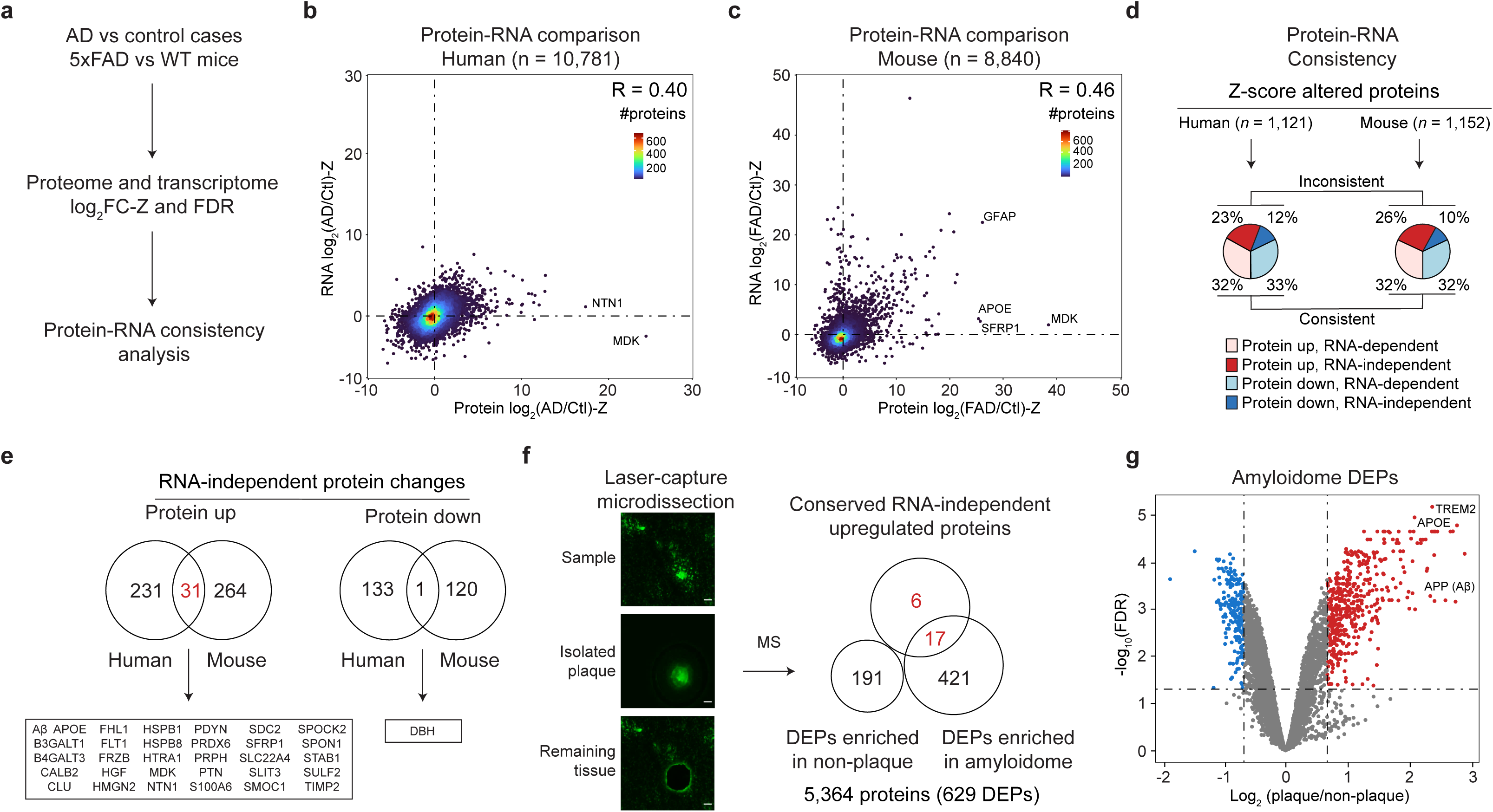
AD and the mouse model show transcriptome-proteome inconsistencies which include RNA-independent upregulated proteins enriched in the amyloidome. **a**, Workflow for comparison of protein/RNA data to define protein-RNA consistencies. **b-c**, Scatterplots of protein-RNA comparisons of log_2_FC-Z in human (*n* = 10,781) and 5xFAD mice (*n* = 8,840). Density is indicated by color gradients. Pearson correlation (R) values are shown. **d**, Percentage of protein-RNA consistency in the population of z-score altered proteins. e, Overlap of RNA-independent protein changes between human and mouse. **f**, Workflow of LCM-MS to compare proteomes in plaque and non-plaque regions, quantifying 5,364 proteins. A Venn diagram illustrated the overlap of 31 shared, RNA-independent, upregulated proteins in both humans and mice with proteins enriched in either plaque or non-plaque regions. **g**, Volcano plot showing proteins enriched in in plaque or non-plaque regions.

We recognized that many of the 31 shared, upregulated proteins are present in amyloid plaques^65–69^, prompting us to fully characterize the amyloidome (i.e. all the components in the amyloid plaque microenvironment) in the 5xFAD mice. We employed laser-capture microdissection (LCM) to isolate amyloid plaques and non-plaque areas from brain tissue (*n* = 4 mice) and profiled the proteome using our modified TMT-LC/LC-MS/MS pipeline, optimized for sub-microgram protein samples^70^. This approach resulted in the quantification of 5,364 proteins (**Fig. 5f-g, Extended Data Fig. 5, Supplementary Table 21**). Quantitative comparison between the plaques and non-plaque areas identified 438 proteins enriched in plaques and 191 in non-plaque areas (FDR < 0.05 and |log_2_FC| > 2SD). Strikingly, of the 31 RNA-independent, upregulated proteins in both human AD and 5xFAD mice, 23 were detected in the amyloidome profiling, in which 17 (74%) were found among the 438 plaque-enriched proteins, while none were detected in the 191 non-plaque-enriched proteins. The results demonstrate that the formation of amyloid plaques contributes to RNA-independent protein accumulation.

### Delayed protein degradation in amyloidome contributes to transcriptome-proteome inconsistency

We hypothesized that proteome-transcriptome discrepancy in AD could be due to reduced protein turnover within the amyloidome. To test this hypothesis, we employed pulsed SILAC labeling (pSILAC)^71–73^ to accurately measure protein turnover rates at high throughput. The 5xFAD mice and WT littermates were fed with heavy lysine SILAC food in a time course (0, 4, 8, 16 and 32 days, with 3 replicates, totaling 30 mice), followed by brain tissue collection and MS profiling (**Fig. 6a, Extended Data Fig. 6**). The kinetics of heavy lysine labeling enabled the determination of protein degradation rates, indicated by protein half-life (T_50_). Apparent T_50_ values were calculated directly from fitting a degradation curve; to account for the recycling of heavy lysine in the mice (**Extended Data Fig. 7**), we derived corrected T_50_ values using an ordinary differential equation model in the JUMPt software^73^.

**Figure 6.**
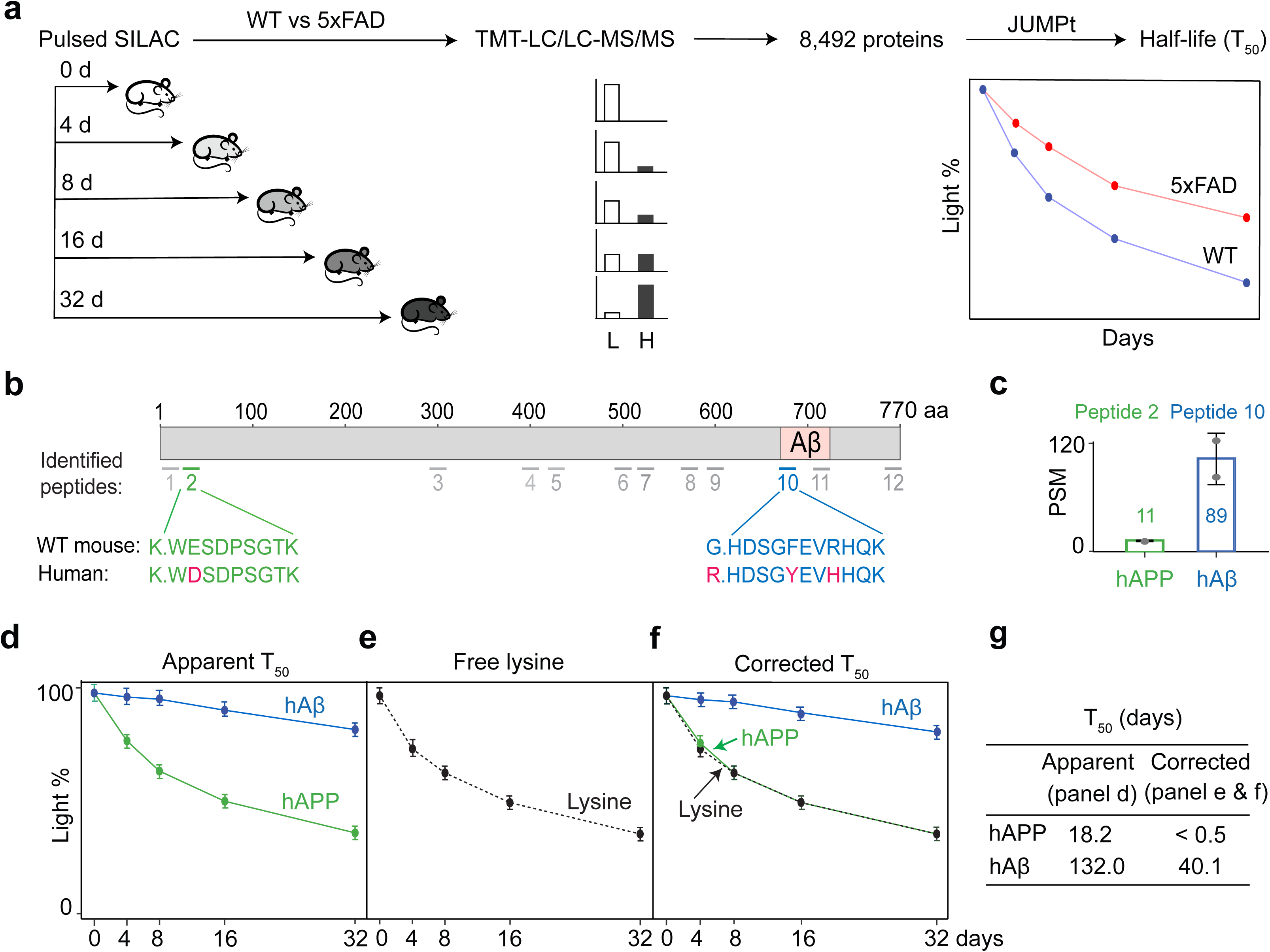
The analysis of AD mouse proteome turnover confirms distinctly different turnover rates for human APP full-length protein and Aβ peptides. **a,** Whole proteome turnover analysis in 5xFAD and WT mice was performed using pulsed SILAC labeling (∼9-month-old, 5 data points, 3 replicates, totaling 30 mice), TMT-LC/LC-MS/MS (2 batches), and the JUMPt program. The analysis covered a comprehensive set of 8,492 unique proteins. **b**, Diagram illustrating the 12 identified peptides in the human and mouse APP or Aβ regions. **c**, PSM counts for the hAPP-specific peptide (peptide 2) and the hAβ surrogate peptide (peptide 10). **d**, Apparent T_50_ values were directly determined from turnover curves for the hAPP- or hAβ-specific peptides. **e,** The curve of free Lys amino acid. **f**, Corrected T_50_ values were calculated based on the distance between the protein curve and free Lys curve, using the JUMPt program, which incorporates a mathematical model to account for delays caused by Lys recycling. **g**. Summary table of hAPP and hAβ T_50_ values. The corrected T_50_ values were much smaller than the apparent T_50_ values.

We quantified 12 tryptic peptides from human APP (hAPP) and mouse APP (mAPP) proteins, both present in 5xFAD (**Fig. 6b, Supplementary Table 22**).Two peptides were human-specific: peptide 2 in the non-Aβ region (used to quantify full length hAPP), and peptide 10 in the Aβ region (used to quantify human Aβ (hAβ) as previous reported)^36^. The peptide spectral match (PSM) counts, a semi-quantitative index^74^, for peptide 10 were significantly higher than those for peptide 2, consistent with the accumulation of Aβ in 5xFAD (**Fig. 6c**). Indeed, hAβ displayed a much longer half-life than hAPP (**Fig. 6d-g**), when analyzing apparent T_50_ (132.0 d for hAβ vs 40.1 d for hAPP) or corrected T_50_ (18.2 d for hAβ vs <0.5 d for hAPP). The results clearly indicate a significantly delayed turnover rate of hAβ relative to hAPP in the 5xFAD mice.

We then analyzed the proteome turnover for the 5xFAD and WT mice, calculating corrected T_50_ values for 8,492 proteins (**Fig. 7a, Supplementary Table 23**). The global T_50_ distributions between 5xFAD and WT were similar with average values around 4-5 d (**Fig. 7b**). A statistical analysis identified 84 proteins with significant changes in half-life between the two genotypes (**Fig. 7c**). 25% of those proteins, including DPP10 and AAK1, exhibited shorter T_50_ in 5xFAD than in WT (**Fig. 7d**), whereas the remaining 75% displayed longer T_50_ in 5xFAD. For example, APOE and VTN showed ΔT_50_ of 4.2 and 3.5 days, respectively (**Fig. 7e**).

**Figure 7.**
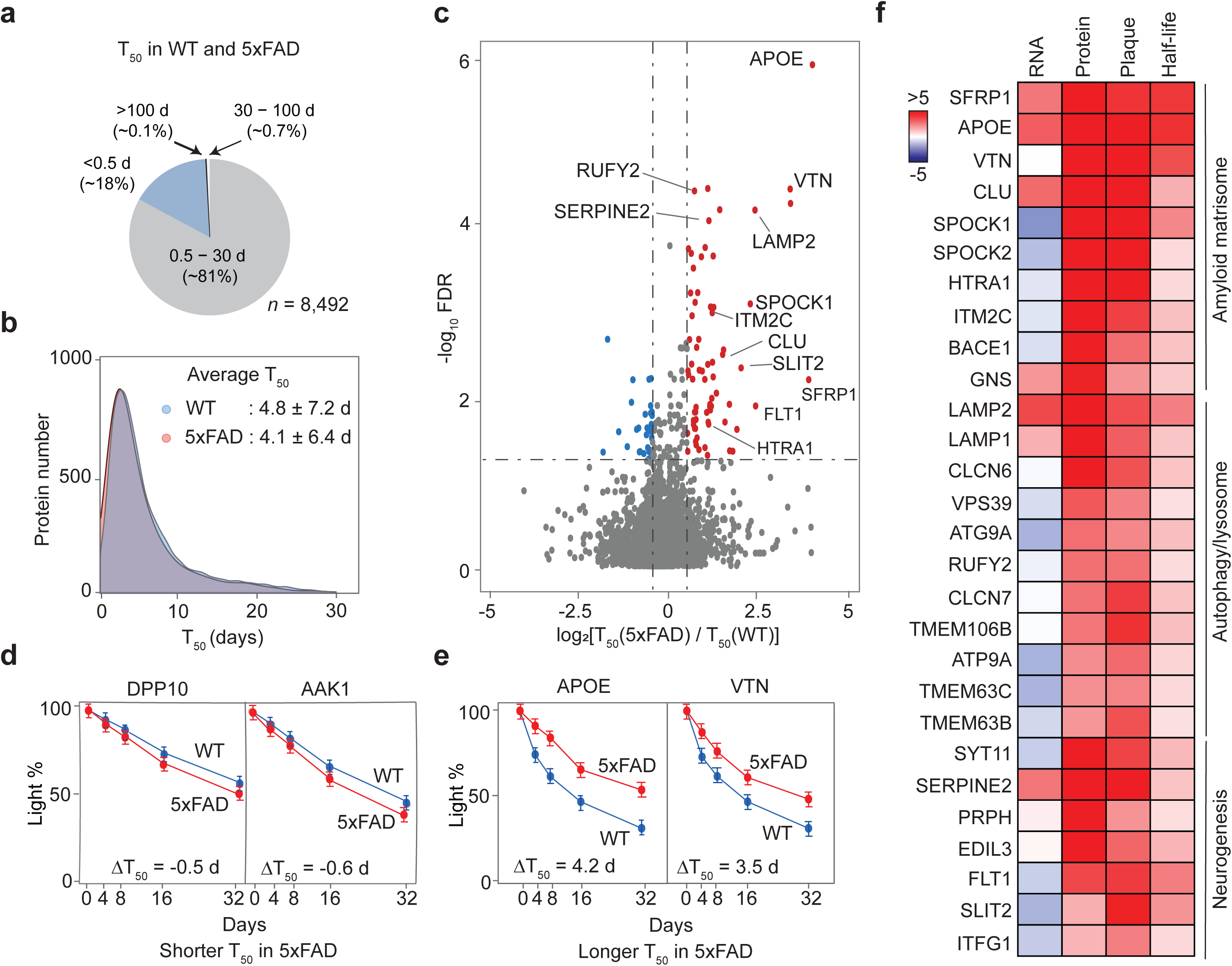
The analysis of AD mouse proteome turnover reveals slow half-lives in amyloidome proteins. **a**, Pie chart displaying the average proportions of corrected protein T_50_ values categorized as very short (<0.5 days), intermediate (0.5-30 days), long (30-100 days), and very long (>100 days) for WT and 5xFAD mice. **b**, Distribution graphs of T_50_ values in both genotypes, showing the average values and standard deviations. **c**, Volcano plots of the log_2_ fold change and FDR for T_50_ in 5xFAD compared to WT, with proteins exhibiting changed T_50_ highlighted in colors and thresholds marked by dashed lines. **d-e**, Examples of proteins that have shortened or extended T_50_ in 5xFAD. **f**, Heatmap illustrating how some proteins with longer T_50_ may be explained by their localization in plaques, contributing to RNA-protein discrepancies. The side bar indicates log_2_FC-Z values for the first three columns or log_2_FC values for the last column.

By integrating transcriptome, proteome, amyloidome, and protein half-lives data from 5xFAD, we found that 32 RNA-independent, upregulated proteins found in the amyloidome showed prolonged half-lives (**Fig. 7f, Supplementary Table 24**). The 32 proteins were enriched in the pathways of amyloid matrisome, autophagy/lysosome and neurogenesis. These findings suggest that the AD transcriptome-proteome discrepancy can be attributed, at least partially, to reduced protein turnover in the amyloidome (**Extended Data Fig. 8**).

## Discussion

### A comprehensive multi-layered proteomics resource for AD mouse models

In this extensive proteomic resource, we have generated the most comprehensive AD mouse brain proteomes to date, analyzing a total of 103 mice across 5 AD models (5xFAD^25,26^, NLF^28^, NLGF^28^, 3xTG^27^, and BiG^45^; **Supplementary Table 2**). This resource also includes phosphoproteome profiling from 36 mice (**Supplementary Table 11**), an in-depth amyloidome analysis from 8 mouse samples (**Supplementary Table 21**), and proteome turnover data from 30 mice (**Supplementary Table 23**). The proteome coverage is high, with most datasets surpassing 10,000 proteins, largely due to our development and implementation of a fully optimized TMT-LC/LC-MS/MS pipeline^55,70,75,76^, extensive fractionation (e.g., at least 40 LC fractions per TMT batch)^46^, and substantial instrument time investment (e.g., ∼4 days per TMT batch). This high-quality proteomics data serve as a critical resource for comparing different mouse models, aligning mouse findings with human AD data, integrating multi-omics datasets, and identifying potential disease-related proteins.

The next-generation of knock-in mouse models for amyloidosis are often considered to have higher physiological relevance and reduced overexpression artifacts compared to 5xFAD^21,23,28,77,78^. Nevertheless, our proteomic comparison highlights similarities between these models, in terms of DEPs (1,382 in 5xFAD, 1,142 in NLGF, **Supplementary Table 7**). A substantial portion (610 proteins) of DEPs overlaps between the two models, and the non-overlapping ones show similar trends in protein alterations. Furthermore, both models have a comparable number of shared DEPs with human consensus data^36,37,52,60^ (159 in 5xFAD, 145 in NLGF, **Supplementary Table 14**). The proteomic correlation with human is also similar between the two models (R = 0.32 in 5xFAD, R = 0.23 in NLGF). Notably, NLGF mice carry the APP Arctic mutation (E693G), which produces a mutated Aβ sequence (E22G) that leads to a slightly different Aβ filament structure compared to the WT Aβ filament in 5xFAD^79,80^. This structural difference may contribute to variations in downstream molecular events. The similar proteomic patterns between the two models suggest they can be effectively used to cross-validate molecular mechanisms related to amyloidosis.

The limitations of mouse models in replicating human AD pathology can largely be attributed to inherent species differences^81^. For example, most mouse models do not exhibit significant neuronal loss. Consistent with the understanding that mouse models cannot fully recapitulate the complexity of human AD events^21,23,78^, our proteomic profiling of five mouse models reveals that each model shares < 25% consensus DEPs of human AD data (**Supplementary Table 14**). Collectively, these mouse models cover ∼42% of human DEPs, indicating that each model has unique characteristics and mimics different subsets of AD molecular events. For instance, tau-related pathways are observed only in 3xTG^27^, whereas some splicing and synaptic defects are unique to the BiG model^45^. This proteomics resource serves as a valuable reference for investigating specific pathways using the most appropriate models.

### Molecular insights from multi-omics integration in AD mouse models

Multi-omics integration offers a robust method to evaluate biological systems and reveal molecular insights^82^, given the generally weak RNA-protein correlation, particularly in the brain which consists mainly of postmitotic, non-dividing cells^35^. In our study, both human and 5xFAD mouse brains exhibit modest RNA-protein correlations, with R values under 0.5, and approximately one-third of protein changes occur independently of RNA levels. Considering protein homeostasis is regulated by events such as modifications, localization, and turnover, we expanded our analysis to include phosphoproteome, subproteome (amyloidome with plaque localization), and protein turnover. While protein phosphorylation did not significantly account for the RNA-protein discrepancies, our multi-layered proteomic data (amyloidome and turnover) supports the hypothesis that amyloid plaque formation create a microenvironment where as many as 32 proteins show delayed turnover, promoting their RNA-independent accumulation in the brain (**Supplementary Table 24**).

Amyloid plaques are dynamic and can grow to 50-100 µm in size^83^. Recent spatial omics has revealed that the Aβ core is enveloped by disease-associated microglia, activated astrocytes, and dysfunctional oligodendrocytes, positioned sequentially, implicating an active microenvironment induced by amyloid plaques (**Extended Data Fig. 8**)^84^. Structural analyses have shown diverse Aβ filament architectures in humans and mice^79,80^, suggesting that their pathological roles are influenced by Aβ-associated proteins within the matrisome^85,86^.

Apoe is a prominent protein in the amyloidome with delayed turnover. It has a well-established role in the “ApoE cascade hypothesis” supported by extensive genetic and biochemical evidence^87^. Primarily produced by astrocytes and also by microglia and neurons^88^, Apoe RNA is upregulated (log_2_FC-Z of 3.26), but its protein change is more dramatic (log_2_FC-Z of 25.38). Apoe shows rapid turnover in WT mice, with a half-life of <0.5 d, but this rate slows significantly to 8.85 d in 5xFAD (**Supplementary Table 23**). These findings indicate that the abundant accumulation of Apoe is driven not only by RNA upregulation but also by changes in protein turnover, possibly due to its direct interaction with Aβ in the plaque microenvironment^87^.

The 32-protein list also includes several understudied proteins, such as Spock1 and Spock2, members of the Sparc proteoglycan family potentially involved in synaptic plasticity^89^; SFRP1, which may promote plaque pathology by inhibiting ADAM10 α-secretase activity in the non-amyloidogenic pathway^90^; and HTRA1, a protease potentially regulating the aggregation and clearance of amyloid proteins^91^. Genetic variations in *HTRA1* gene are linked to age-related macular degeneration^92,93^. These proteins are also found on the upregulated consensus protein list in AD brains (**Supplementary Table 23**), implicating a possible role in the formation of amyloid plaque microenvironment.

Interestingly, we found a number of proteins enriched in the autophagy/lysosome pathway with delayed turnover rates (**Supplementary Table 24**), such as Tmem106b, which has been identified as an aggregated filament protein in several neurodegenerative disorders including AD^94–96^, and genetically linked to frontotemporal lobar degeneration (FTLD)^97^. Tmem106b accumulates in lysosomes, playing a role in lysosomal dysfunction^98^. The slow turnover of lysosomal proteins in AD mice might indicate defective autophagic degradation, possibly because lysosomes are damaged by intracellular Aβ species^99,100^. This disruption could impair cellular homeostasis, and further inhibition of lysosomal function exacerbates AD-related phenotypes, underscoring the pivotal role of lysosomal pathways in disease progression.

In summary, our study presents a comprehensive multi-layered proteomics resource, profiling five AD mouse models and providing insights into proteomic responses to AD pathologies. Through whole proteome analysis, along with phosphoproteome, amyloidome, and turnover data, the resource supports the hypothesis that amyloid plaques create a microenvironment that promotes protein accumulation. Compared with human proteomics data, this resource enables researchers to select relevant disease pathways for study in appropriate mouse models, with the potential to develop other AD models in the future. All data from this resource is freely accessible on a website (https://penglab.shinyapps.io/mouse_ad_profile/).

## Methods

### Mouse models in proteome analysis

The 5xFAD transgenic mice^25,26^ and 3xTG mice^27^ were purchased from The Jackson Laboratory (stock #034848 and #034830, respectively), while the NLF and NLGF KI mice^28^ were provided by Dr. Takaomi Saido at the RIKEN Center for Brain Science. The BiG mice were generated by crossing 5xFAD with N40K transgenic mice as described previously^45^. These mice were maintained in the Animal Resource Center at St. Jude Children’s Research Hospital or the University of Arizona according to the Guidelines for the Care and Use of Laboratory Animals. All animal procedures were approved by the Institutional Animal Care and Use Committee (IACUC). The brain tissues were collected at various ages, rapidly dissected, and then immediately frozen on dry ice before being stored at -80 °C.

### Mouse SILAC labeling for protein turnover analysis

The mice were labeled using Mouse Express® L-Lysine (^13^C_6_, 99%) Mouse Feed (5 g per day, Cambridge Isotopes Laboratories). The mice were conditioned by providing the light SILAC food for 3 d before labeling, and then fed with the heavy SILAC food in a time course. Cortical brain tissue samples were harvested for turnover analysis. The fully labeled mice were generated as previously reported^101^. The heavy mouse chow was used to feed wild type mice from the parental generation through to the F2 generation. Through two generations, the mouse proteins were fully labeled^101^.

### Collection of amyloid plaques by LCM

LCM was performed essentially according to a previously reported method^65^. Mouse brain was embedded in OCT Compound (Jed Pella Inc., Redding, CA), sectioned at 12 μm in a cryostat and mounted on Arcturus Pen membrane glass slides (LCM0522, ThermoFisher). The sections were thawed, fixed with 75% ethanol for 1 min, stained with 1% thioflavin-S (MilliporeSigma) or X-34 (MilliporeSigma) for 1 min, washed in 75% ethanol for 1 min, dehydrated, cleared in xylene and air-dried. LCM was performed using an Arcturus XT Laser Capture Microdissection System (Arcturus, ThermoFisher) with the following settings: excitation wavelength, 495 nm; laser power, 60–80 milliwatts; duration, 1 ms; laser spot size, 7.5 µm, and 30 µm collection diameter. About 500 amyloid plaques were procured from each section, while non-plaque areas were captured as a control. The captured samples were stored at -80 °C.

### Protein profiling by TMT-LC/LC-MS/MS analysis

The experiments were performed according to our previously optimized protocol^102,103^. Briefly, the mouse brain samples were weighed and homogenized in lysis buffer (8 M urea, 50 mM HEPES, pH 8.5, and 0.5% sodium deoxycholate,100 µL buffer and ∼20 µL beads per 10 mg tissue) with 1 x PhosSTOP phosphatase inhibitor cocktail (Roche). ∼50 µg protein from each sample was then digested in two steps by Lys-C and trypsin, with DTT reduction and iodoacetamide alkylation, followed by desalting with a C18 Ultra-Micro SpinColumn (Harvard apparatus). The desalted peptides were resuspended in 50 mM HEPES (pH 8.5) to a concentration of ∼1 µg/µL, and fully labeled with TMT or TMTpro reagents. The reaction was quenched, equally pooled, and desalted for the subsequent prefractionation.

The pooled TMT samples were fractionated by offline basic reverse phase (RP) LC with an XBridge C18 column (3.5 μm particle size, 4.6 mm x 25 cm, Waters; buffer A: 10 mM ammonium formate in H_2_O, pH 8.0; buffer B: 10 mM ammonium formate in 90% acetonitrile, pH 8.0). Fractions were collected in a gradient of 15-42% buffer B, and then concatenated into at least 40 samples to maintain high-resolution power. The concatenated samples were dried by SpeedVac, resuspended in 5% formic acid (FA), and analyzed by Q-Exactive HF Orbitrap MS (Thermo Fisher Scientific) in a 95 min nano-LC gradient of 15%-48% buffer B (buffer A: 0.2% FA, 5% DMSO; buffer B: buffer A plus 65% acetonitrile). MS1 scan settings were 60,000 resolution, 410-1600 *m/z* scan range, 1 x 10^6^ AGC, and 50 ms maximal ion time. MS2 settings were 20 data-dependent MS2 scans, 60,000 resolutions, starting from 120 *m/z*, 1 x 10^5^ AGC, 120 maximal ion time, 1.0 *m/z* isolation window with 0.2 *m/z* offset, HCD, 32% specified normalized collision energy, and 15 s dynamic exclusion^70^.

### Phosphopeptide enrichment

The basic pH RPLC-fractionated, TMT labeled peptides were concatenated to 10 fractions (∼0.3 mg per fraction), dried, and resuspended in binding buffer (65% acetonitrile, 2% TFA, and 1 mM KH_2_PO_4_). TiO_2_ beads (0.9 mg per sample, GL sciences) were incubated with the peptide fraction at 21°C for 20 min. The TiO_2_ beads were then washed twice with washing buffer (65% acetonitrile, 0.1% TFA) and packed into a C18 StageTip (Thermo Fisher), followed by phosphopeptide elution with the basic pH buffer (15% NH_4_OH, and 40% acetonitrile). The eluates were dried and dissolved in 5% formic acid for LC-MS/MS analysis^55^.

### Protein identification and quantitation in the TMT analysis

The protein identification and quantification were analyzed using the JUMP software suite^47^. The MS data were searched against the protein database merged from Swiss-Prot, TrEMBL (from Uniprot), and UCSC databases (mouse: 59,423 entries). To evaluate the false discovery rate (FDR), decoys were generated by reversing the target protein sequence^104^. Search parameters included precursor ion and product ion mass tolerance (10 ppm), maximal modification sites (n = 3), full trypticity, maximal missed cleavage (n = 2), static modification of TMT tag (+304.20715), methionine oxidation dynamic modification (+15.99491), and cysteine carbamidomethyl static modification (+57.02146) if the residue was alkylated with iodoacetamide. In the pSILAC-TMT analysis, the MS raw files were searched twice with or without Lys labeling (+6.02013). Peptide-spectrum matches (PSMs) were filtered by matching scores and mass accuracy to keep protein FDR below 1%. The peptides shared by multiple homologous proteins were assigned by the software to the protein with the maximal PSM number, based on the rule of parsimony. The protein quantitation was performed using the TMT reporter ion based on a published method^46^. In the case of phosphoproteome profiling, we applied the JUMPl program^36^ with the phosphoRS algorithm^105^ to the analysis of phosphosite localization scores (Lscore, 0%–100%) for each PSM, and then determine the appropriate phosphosites.

### Differential expression analysis

Proteomic analysis was performed on mouse brain samples, followed by log_2_ transformation and median normalization. Principal component analysis was used to detect outliers and evaluate the effect of variables such as batch bias, sex, age and genotype, with the R package *prcomp*^106^. DE analysis was performed via a moderated *t*-test from the *MKmisc* package^107^. The *p* values were adjusted to FDR using the Benjamini-Hochberg procedure^108^. DEPs were accepted if they passed the FDR cutoff (0.05) and a log_2_FC cutoff (2 SD).

### Pathway enrichment and protein-protein interaction (PPI) network analysis

Pathway enrichment of DEPs was performed by GO enrichment analysis^50^ and further analyzed by the PANTHER^109^ overrepresentation test (Fisher’s Exact test). Results were filtered by FDR to identify DEP-associated pathways. DEPs within the pathways were superimposed against a custom PPI database, which combined the InWeb_IM^110^, STRING^111^, and BioPlex^112^ databases, as detailed previously^36^. Interacting DEPs were then visualized in Cytoscape^113^.

### Differentially altered kinase activity inference by KSEA

The analysis was performed using the ‘KSEAapp’ R package (v0.99.0) within RStudio (v4.1.2), with 122 DE phosphopeptides as input^59^. Both PhosphoSitePlus^114^ and NetworKIN^115^ databases were utilized to find kinase-substrate interactions and phosphosite information. Subsequently, kinase substrates were extracted to derive the corresponding kinase activities. For these kinases, redundancy caused by shared substrates was manually curated. The log_2_FC values, representing the mean log_2_FC of all the kinase’s substrates, were used to generate the heatmap.

### Summarization of pathway activities from individual components

We used a previously modified pathway activity inference strategy^61,62^ to derive the activity of a given pathway, termed a(P), in AD and mouse model samples:

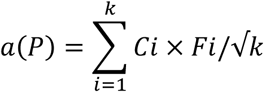

Where *k* represents the number of proteins in each pathway, *F_i_* is the Log_2_FC for individual proteins, and *C_i_* denotes the functional annotation of the protein, assigned as either +1 for proteins with an activation role or -1 for proteins with an inhibitory role.

### Prioritization of proteins/genes based on multi-omics datasets

We implemented a gene/protein ranking method based on order statistics for multi-omics integration^36^. This approach combines *N* distinct protein/gene ranking sets into a single comprehensive ranking. In this analysis, we utilized a total of 12 individual datasets: GWAS-identified risk loci^13–20,36^, human transcriptome^116^, MCI proteome^36^, AD proteome datasets^36,37^, AD aggregated proteome^117^, AD phosphoproteome^36^, 5xFAD transcriptome^36^, AD amyloidosis proteome integrated from our 5xFAD and NLGF data, 3xTG proteome, BiG proteome, AD amyloidosis phosphoproteome, and interactome closeness to known AD genes by PPI network distance^36^.

### RNA-protein consistency analysis

To account for scale difference, proteomics and transcriptomics^36^ data were converted to Z scores, generating log_2_FC-Z data. RNAs and proteins with shared accessions were used for comparison. Proteins were grouped as upregulated (Z > 2) or downregulated (Z < -2). Protein-RNA consistency was determined by a ΔZ (the Z score difference between protein and RNA) absolute value larger than 2.5. However, if both RNA and protein had absolute Z-values larger than 4 and changed in the same direction, the pair was still considered to be consistent, regardless of ΔZ.

### Pulse SILAC-TMT hyperplexing data processing

Following protein identification and quantification using the JUMP software suite^47^, the MS raw data and JUMP output files were further processed to remove TMT noise for accurate quantification, utilizing the JUMPsilactmt Python program (version 1.0.0). Briefly, noise levels were identified in light PSMs (the fully labeled SILAC channel) and heavy PSMs (the unlabeled channel) and subtracted from other channels (see **Extended Data Fig. 6** for details). However, the TMT intensities from adjacent light/heavy PSM pairs could not be directly compared because they were produced from distinct MS scans. To address this, the JUMPsilactmt quantified the composite MS1 heavy and light ion intensities as in the SILAC quantification method^73^, and used their ratio to normalize the denoised MS2 TMT reporter ion intensities in different MS scans. The normalized heavy and light TMT intensities were converted into a fraction of light (L%). Finally, quantification at the PSM level was summed to the peptide level and subsequently to the protein level. L% values were averaged across biological replicates for the JUMPt analysis.

### Protein turnover rate analysis by JUMPt

Global protein half-lives were determined using JUMPt software^73^ (version 1.0.0). First, we calculated the apparent T_50_ for each protein individually, using the averaged L% of the protein across time points, without accounting for in vivo Lys recycling. Next, we analyzed the corrected T50 under “Setting-2,” incorporating free Lys data and L% to estimate all protein T_50_ values simultaneously using an ordinary differential equation model. It should be noted that, for proteins with very short T_50_ (< 0.5 days) or very long T_50_ (> 100 days), the calculation of corrected T_50_ analysis is less reliable, though only a small fraction of proteins exhibits these extreme half-lives.

### Analysis of mouse protein turnover changes

To identify statistically significant changes in T_50_ between different genotypes, the L% of each protein over the time course was analyzed by two-way ANOVA, with genotype (WT vs. 5xFAD) and labeling time (4, 8, 16, and 32 d) as variables. The ANOVA *p* values for genotypes were then adjusted to FDR by the Benjamini-Hochberg procedure^108^. Moreover, the ΔT_50_ for each protein was calculated as the logarithm (base 2) over the corrected T_50_ ratio between 5xFAD and WT mice. The proteins were filtered by FDR (<0.05) and |log_2_FC| ( 2 SD) to generate the final list with altered half-lives.

## Data availability

RAW data results have been deposited in the PRIDE database (https://www.proteomexchange.org) with accession numbers PXD018590, PXD007974, PXD023395, PXD031545, PXD031732, PXD031734, PXD031735, PXD031769, and PXD053314 (**Supplementary Table 2**). The program of JUMPsilactmt (version 1.0.0) is publicly available from GitHub (https://github.com/abhijitju06/JUMPsilactmt-Version-0.0.1). The program of JUMPt (version 1.0.0) is available from GitHub (https://github.com/abhijitju06/JUMPt-Version-1.0.0). The program details, along with example input and output files, are provided on the respective websites.

## Acknowledgements

We thank St. Jude Shared Resources and Core Facilities, including Animal Research Center, Transgenic/Gene Knockout, Proteomics and Metabolomics. We thank Takaomi C. Saido and RIKEN Brain Science Institute for providing APP^NLF^ and APP^NLGF^ mice. We thank Ines Chen for critical reading and editing. This work was partially supported by National Institutes of Health grants R01AG047928 (J.P.), R01AG053987 (J.P.), RF1AG068581 (J.P.), U54NS110435 (J.P.), U19AG069701 (J.P.), RF1AG064909 (G.Y. and J.P.), and the ALSAC foundation.

## Contributions

J.P., X.H., G.Y. and Y.L. conceived this project. J.M.Y., X.H., K.Yang, D.L., M.Z., Z.W., K.Yu, D.G.L., D.V., M.N., H.S, B.X., P.-C.C, Y.J., X.Z., Z.W., B.V., Q.H., A.T., P.T.R., and R.C. performed the experiments. J.M.Y., X.H., A.D., K.Yang, H.K.S., Y.F., Y.L., Z.-F.Y., X.W., S.P., G.Y., Y.L. and J.P. analyzed the data. J.M.Y., X.H., A.D., and J.P. wrote the manuscript.

---

**Extended Data Fig. 1.**
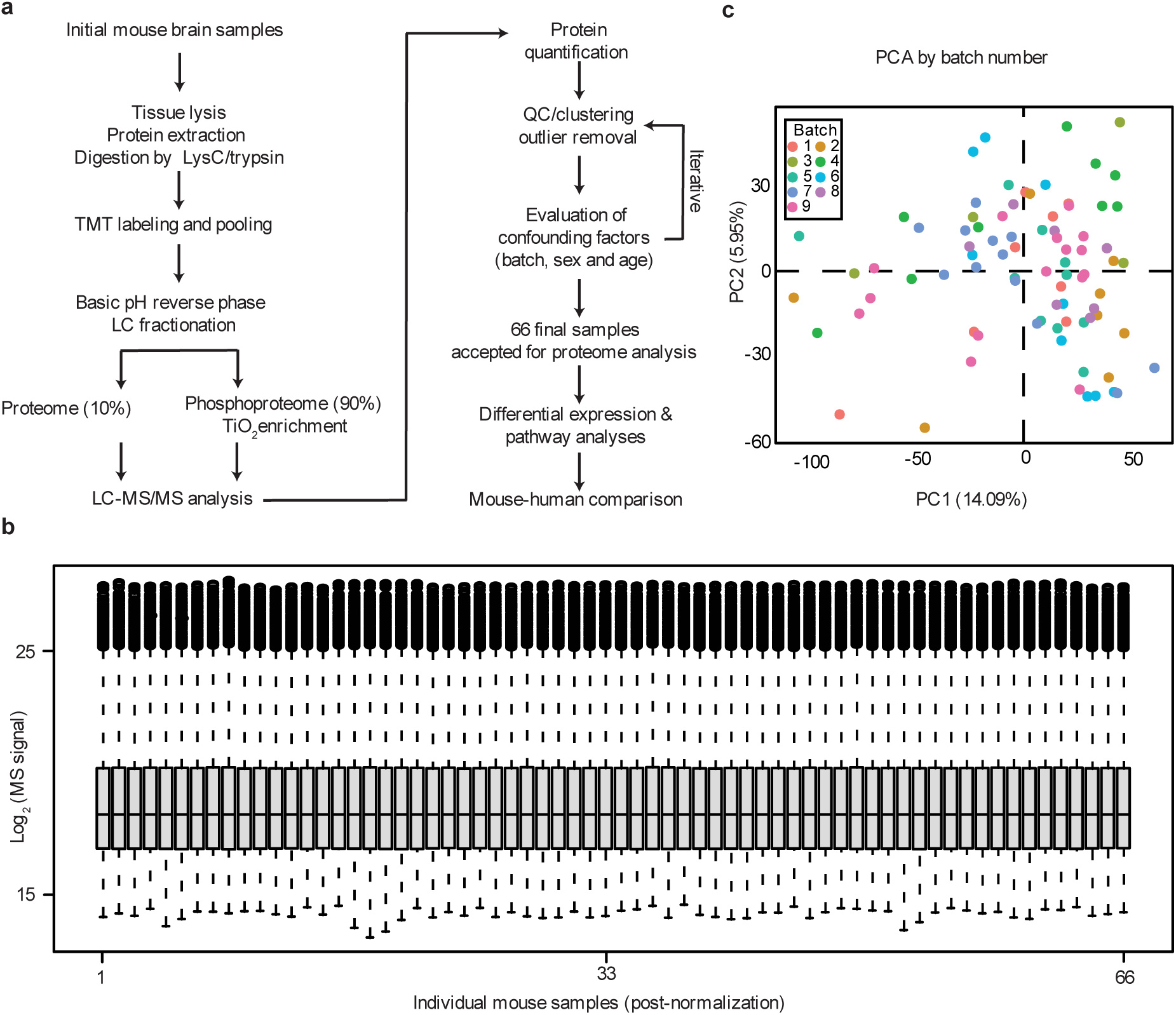
Workflow and quality control of mouse proteomic samples. **a**, TMT-LC/LC-MS/MS and quality control workflow for analysis. Finally, 66 mouse samples were used in our proteome analysis. **b**, Box plot to show the distribution of normalized protein intensities in the mouse samples. **c**, Principal component analysis of mouse samples in different TMT batches. No batch bias was observed.

**Extended Data Fig. 2.**
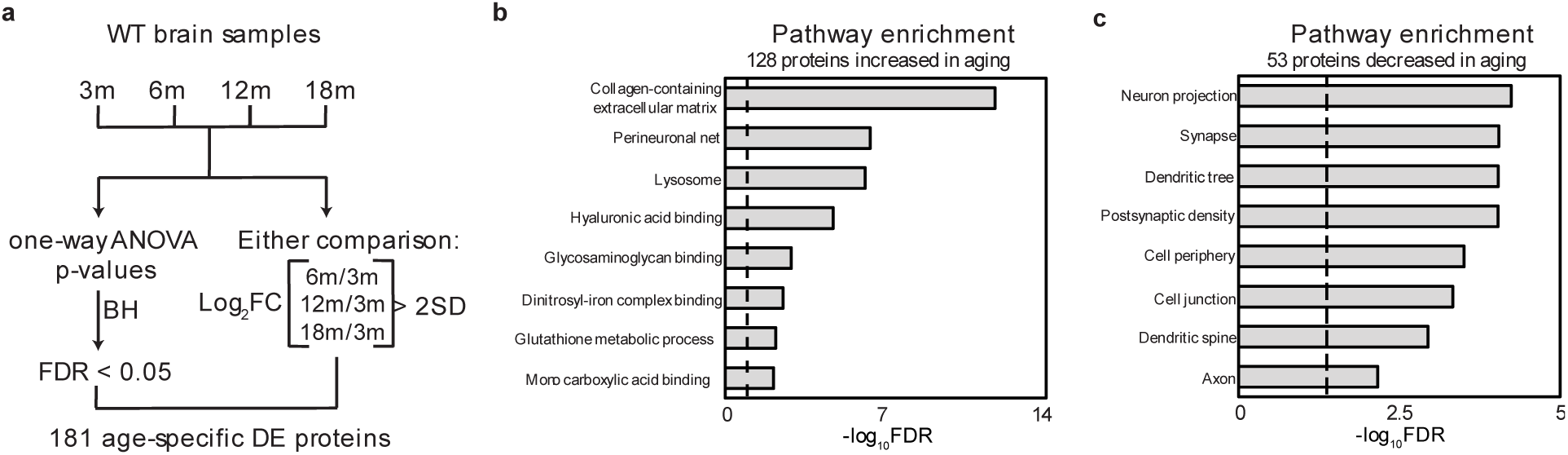
Proteomic analysis of age-linked alterations in WT mice. **a**, Workflow of age-linked proteomics analysis. **b**, The cluster of upregulated proteins with age. Each line represents one protein. **c**, The cluster of downregulated proteins with age.

**Extended Data Fig. 3.**
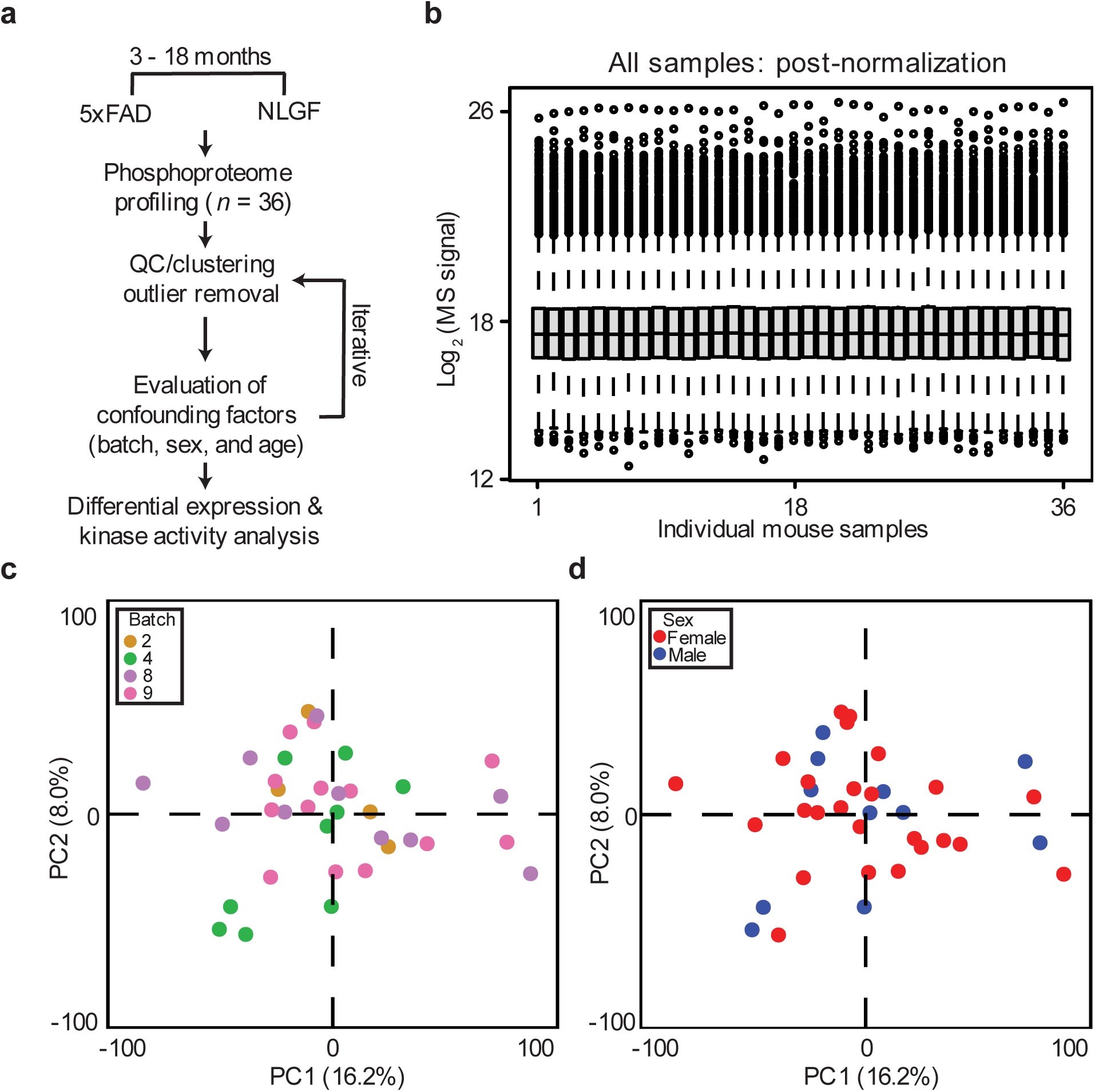
Phosphoproteomic and quality control analysis of 5xFAD, NLGF and WT mice. **a**, Workflow for phosphoproteomics analysis. **b**, Distribution of measured phosphoprotein intensities in mouse samples. **c**, Principal component analysis of mouse samples highlighting the normalization of batch. **d**, Principal component analysis of mouse samples showing no obvious sex effect.

**Extended Data Fig. 4.**
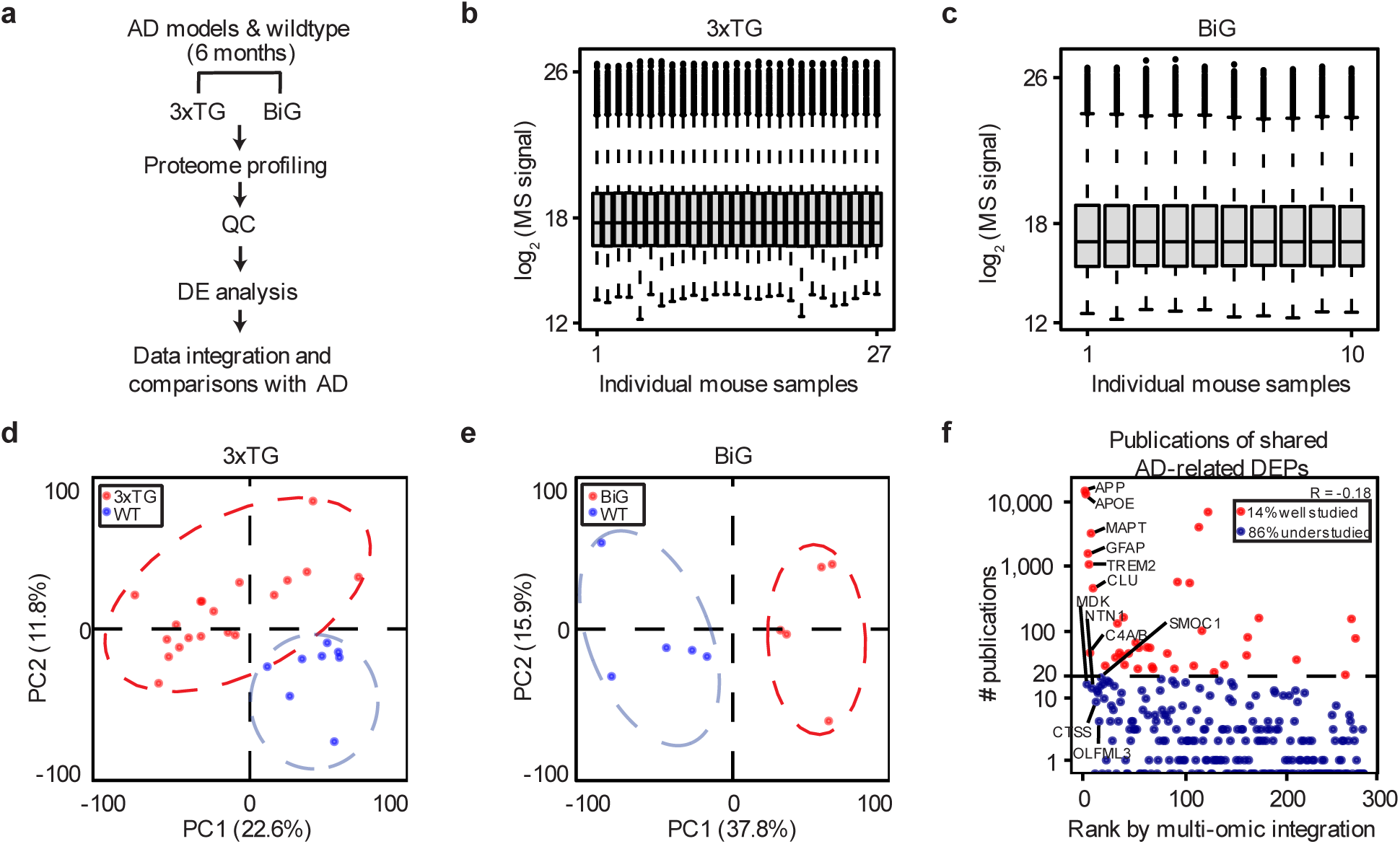
Whole proteome analysis of 3xTG, BiG and WT mice and the publication record of the AD-mouse shared proteins. **a**, Workflow for whole proteome analysis by TMT-LC/LC-MS/MS. **b**, Distribution of normalized protein intensities in 3xTG and WT samples. **c**, Distribution of normalized protein intensities in BiG and WT samples. **d**, Principal component analysis of 3xTG and WT samples highlighting the difference between genotypes. **e**, Principal component analysis of BiG and WT samples highlighting the difference between genotypes. **f**, Publication record for the 275 AD-mouse shared proteins is represented by the number of publications and their integrative ranks determined by multi-omics analysis. The number of publications were found by PubMed search using “Alzheimer” and protein gene names in August 2024. Using 20 publications as the cutoff, 14% of the proteins are well-studied, whereas 86% are under-researched. Notably, Pearson correlation analysis between the rank number and publication count yielded a negative value, suggesting that published research is disproportionately focused on a few well-known AD genes/proteins.

**Extended Data Fig. 5.**
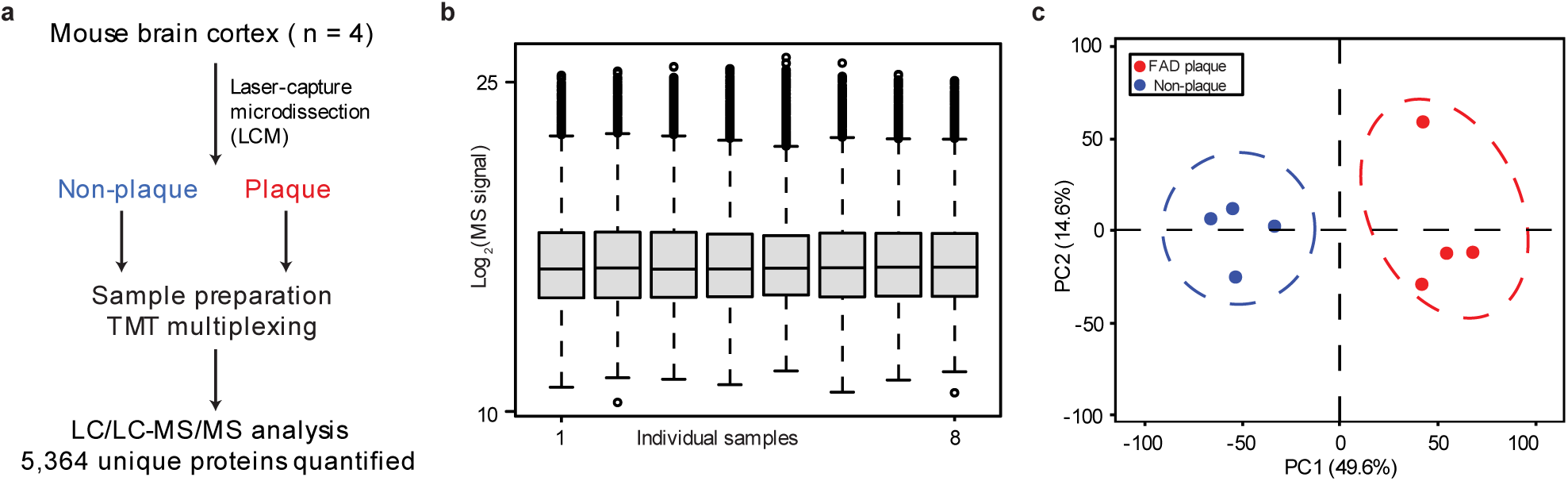
Plaque proteome analysis of AD mice. **a**, Workflow for LCM-TMT-LC/LC-MS/MS. **b**, Distribution of normalized protein intensities in plaque and non-plaque samples. **c**, Principal component analysis highlighting the clear difference between the plaque and non-plaque results.

**Extended Data Fig. 6.**
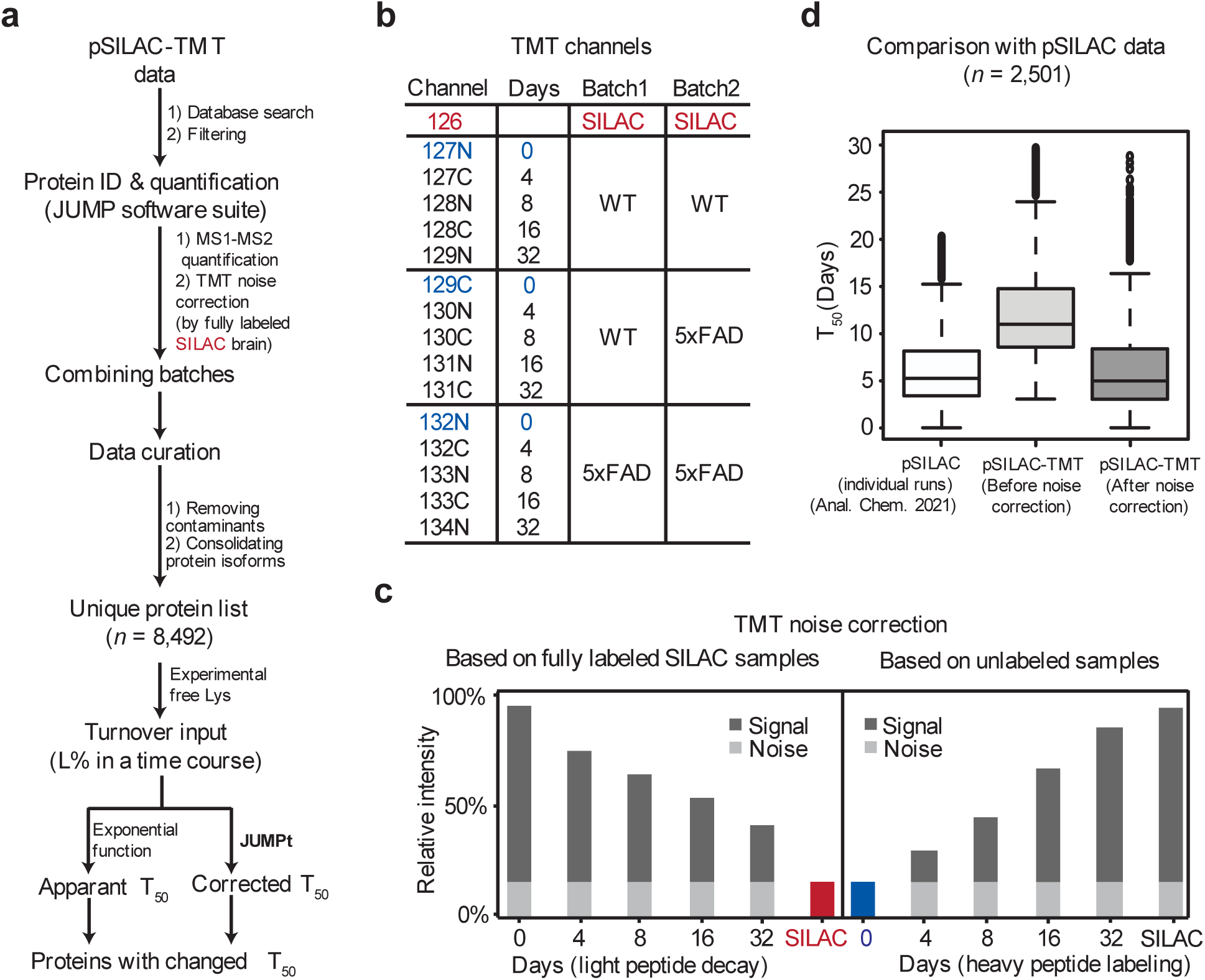
Comprehensive workflow for mouse pSILAC-TMT-LC/LC-MS/MS analysis and the importance of TMT noise correction. **a**, Workflow including mouse pSILAC labeling, TMT-LC/LC-MS/MS experiments, protein identification and quantification via JUMP, batch merging, data curation, and JUMPt analysis for both apparent and corrected T50 turnover rates. **b**, TMT channels for proteomics experiments designed to include three replicates each of WT and 5xFAD mice, with five-time points (0, 4, 8, 16, and 32 days) distributed across two batches. The fully SILAC-labeled mouse brain tissues from two mouse generations^101^ were highlighted in red, serving as a basis for noise correction. **c**, Schematic diagram illustrating the correction of TMT noise. In the light peptide decay curve, the fully labeled SILAC sample (red) facilitated noise detection since all Lys peptides were in the heavy form. Conversely, for the heavy peptide decay curve, the unlabeled sample (blue) from day 0 was utilized to determine the noise level. **d**, Comparison of protein half-lives in WT brain samples using the pSILAC and pSILAC-TMT methods, using 2,501 overlapping proteins. In the pSILAC method, each sample was analyzed separately using the traditional approach, effectively avoiding ratio compression in the dataset ^73^. Initially, before TMT noise correction, the global half-lives of proteins from the pSILAC-TMT experiment were significantly higher compared to those from traditional pSILAC analysis. After implementing TMT noise correction, the half-lives in the two datasets aligned closely, demonstrating that the TMT noise was effectively eliminated.

**Extended Data Fig. 7.**
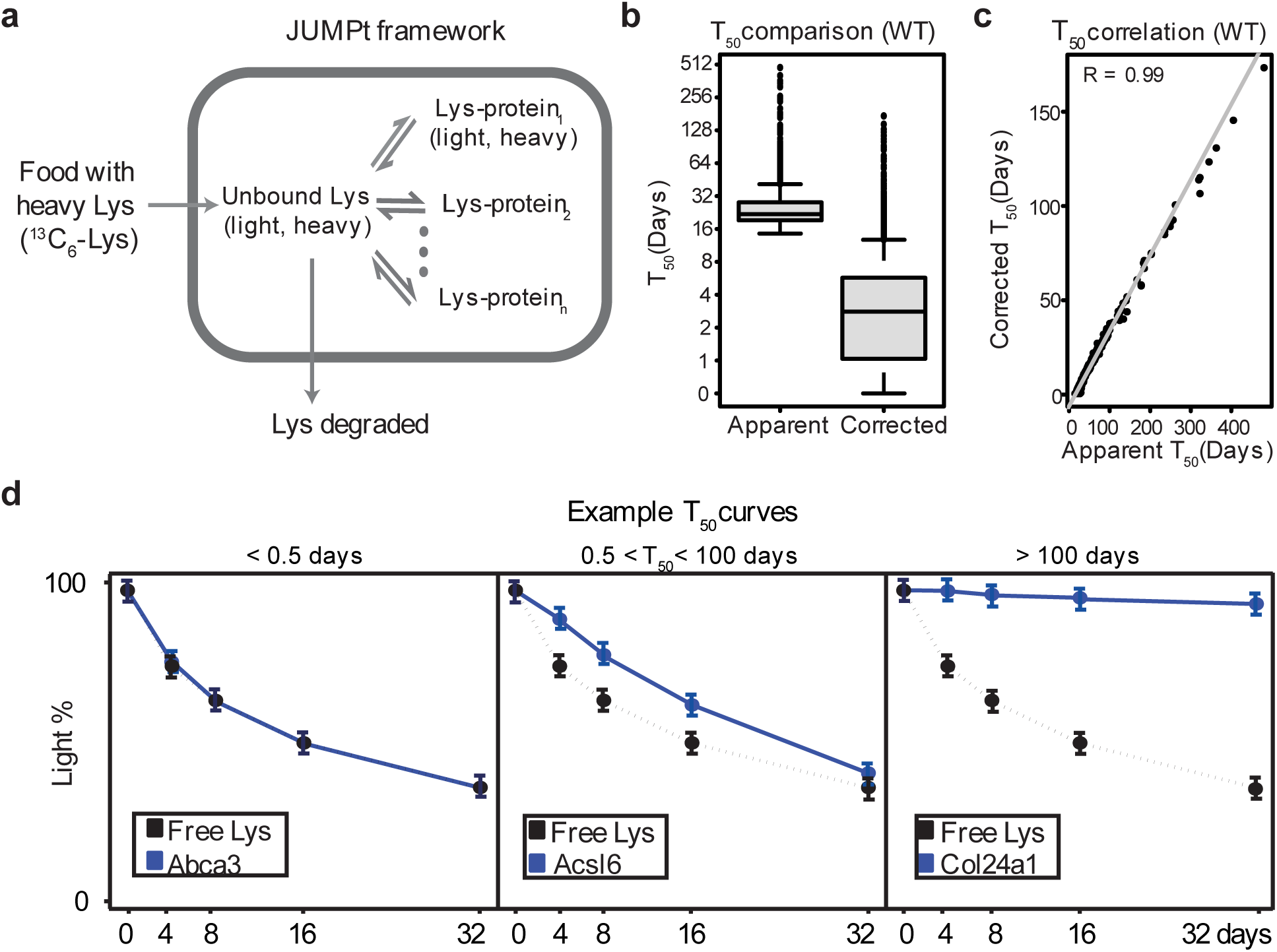
The principle of half-life analysis by JUMPt considering the recycling of Lys in mice. **a**, Kinetic model for the JUMPt program: Heavy Lys is absorbed from food and transported into cells, contributing to a mixed pool containing previous light and newly absorbed heavy Lys. A portion of the Lys in cells is degraded, while the remainder is incorporated into proteins. When proteins degrade, the Lys in proteins is recycled back into the free Lys pool. **b**, The differences between apparent and corrected half-lives. **c**, Correlation between apparent and corrected half-lives. **d**, Examples of proteins with very short, intermediate, and very long half-lives: Proteins with very short half-lives have decay curves nearly overlapping with the free Lys curve, complicating accurate half-life calculation. These half-life values were set below 0.5 day. Conversely, proteins that decay very slowly have extremely long half-lives, presenting challenges for accurate computation. These half-life values were set above 100 days.

**Extended Data Fig. 8.**
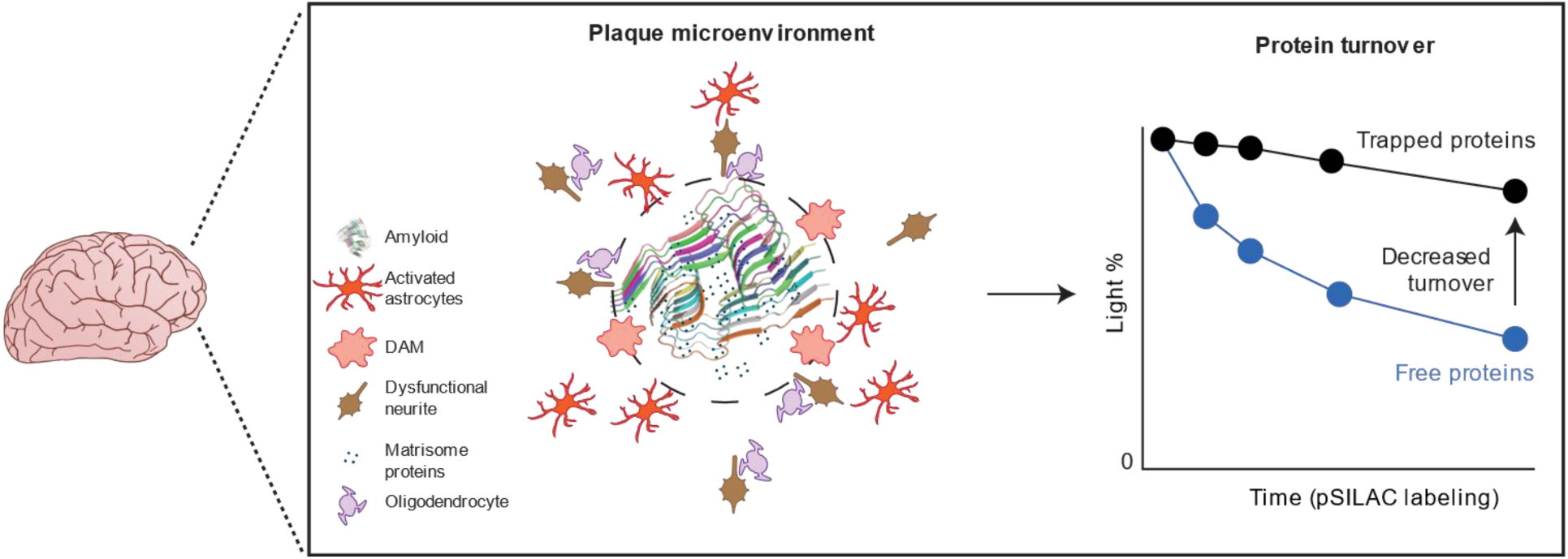
Model showing components plaque microenvironment and the changes in protein turnover in amyloid plaque microenvironments. The core-shell model of plaque microenvironment is adapted from published spatial omics data^84^.

